# Complexity-multistability relationships: How does species diversity shape the number of alternative stable states?

**DOI:** 10.64898/2026.07.30.741147

**Authors:** Gen Iwashita, Shota Shibasaki, Kenta Suzuki, Hirokazu Toju, Masato Yamamichi

## Abstract

Ecologists have long investigated how community complexity affects ecological stability, yet how community complexity influences multistability, defined as the presence of alternative stable states, remains poorly understood. We developed a novel framework integrating stochastic community assembly with stability landscape analysis to quantify multistability from species interaction matrices. Using this framework, we systematically explored how species interaction properties shape the relationship between species diversity (species pool size) and the number of alternative stable states. Mean interaction strength was the primary determinant: competitive interactions amplified the positive relationship between species diversity and the number of alternative stable states. In competitive communities, a greater number of alternative stable states was associated with lower community uncertainty, a measure of the long-term unpredictability of community assembly dynamics. These results highlight the importance of characterizing the entire stability landscape. Our framework provides a general approach for understanding and quantifying multistability in complex ecological communities.

## 1 Introduction

How community complexity influences ecological stability has long been a central question in community ecology (reviewed by Pimm 1984, McCann 2000, Landi et al. 2018). Community complexity encompasses species richness and properties of species interactions, including interaction strength, connectance, and network topology. Classic studies assumed that ecological communities become more stable as species richness increases (MacArthur 1955), but dynamic models revealed that increasing species richness, interaction strength, and connectance can destabilize ecological communities (Gardner and Ashby 1970, May 1972, Allesina and Tang 2012). Subsequent studies showed that the complexity–stability relationship depends on various community-scale characteristics and mechanisms such as mutualism (Mougi and Kondoh 2012), juvenile–adult stage structure (de Roos 2021, Giménez-Romero et al. 2025), nonlinear density dependence (Hatton et al. 2024), spatial structure (Gravel et al. 2016, Mougi and Kondoh 2016), and interaction network topology (Okuyama and Holland 2008, Grilli et al. 2016). Ecological stability in these studies has typically been quantified using resilience, resistance, robustness, and other related metrics that characterize community responses to perturbations around a single stable state (Kéfi et al. 2019). By contrast, multistability describes the existence of multiple stable states under identical environmental conditions, yet the effects of community complexity on multistability remain poorly understood (Beisner et al. 2003, Kéfi et al. 2016).

Multistability has profound consequences for community dynamics. Shallow lakes provide a classic example, with a clear-water state dominated by submerged macrophytes and a turbid state dominated by phytoplankton (Scheffer 2004). These alternative stable states are maintained by positive feedbacks involving light and nutrient cycling (Kéfi et al. 2016). When multistability exists, environmental conditions alone are insufficient to predict long-term community dynamics because identical environmental conditions can support different stable states. Consequently, realized community states depend on historical processes, including community assembly history and previous perturbations. This historical contingency gives rise to priority effects during community assembly (Fukami 2015, Ke and Letten 2018). Moreover, even small environmental perturbations can trigger abrupt regime shifts between stable states, leading to hysteresis. Numerous empirical and theoretical studies have demonstrated such regime shifts and hysteresis across ecosystems, including coral reefs, grasslands, and forests/savannas (Scheffer et al. 2001, Scheffer and Carpenter 2003, Schröder et al. 2005). The possibility of abrupt regime shifts has motivated the development of early warning signals for critical transitions (Scheffer et al. 2012, Scheffer et al. 2015).

Despite its ecological importance, a general understanding of how community complexity shapes multistability remains lacking. This is partly because identifying and quantifying alternative stable states remains computationally challenging, particularly in species-rich communities. Several theoretical studies have suggested that the number of alternative stable states often increases with species richness (Fig. 1a; Gilpin and Case 1976, van Nes and Scheffer 2004, Yamamichi et al. 2014, Fried et al. 2016, Altieri et al. 2021, Ros et al. 2023, Aguadé-Gorgorió et al. 2024). However, the diversity–multistability relationship depends strongly on species interaction properties. For example, the number of alternative stable states shows a concave relationship with species richness under strong and symmetric competition (Fried et al. 2016), whereas it increases exponentially with species richness under symmetric random interactions (Altieri et al. 2021). Nevertheless, a general framework for quantifying multistability across complex ecological communities is still lacking, limiting our ability to evaluate how species interaction properties shape it.

**Figure 1.**
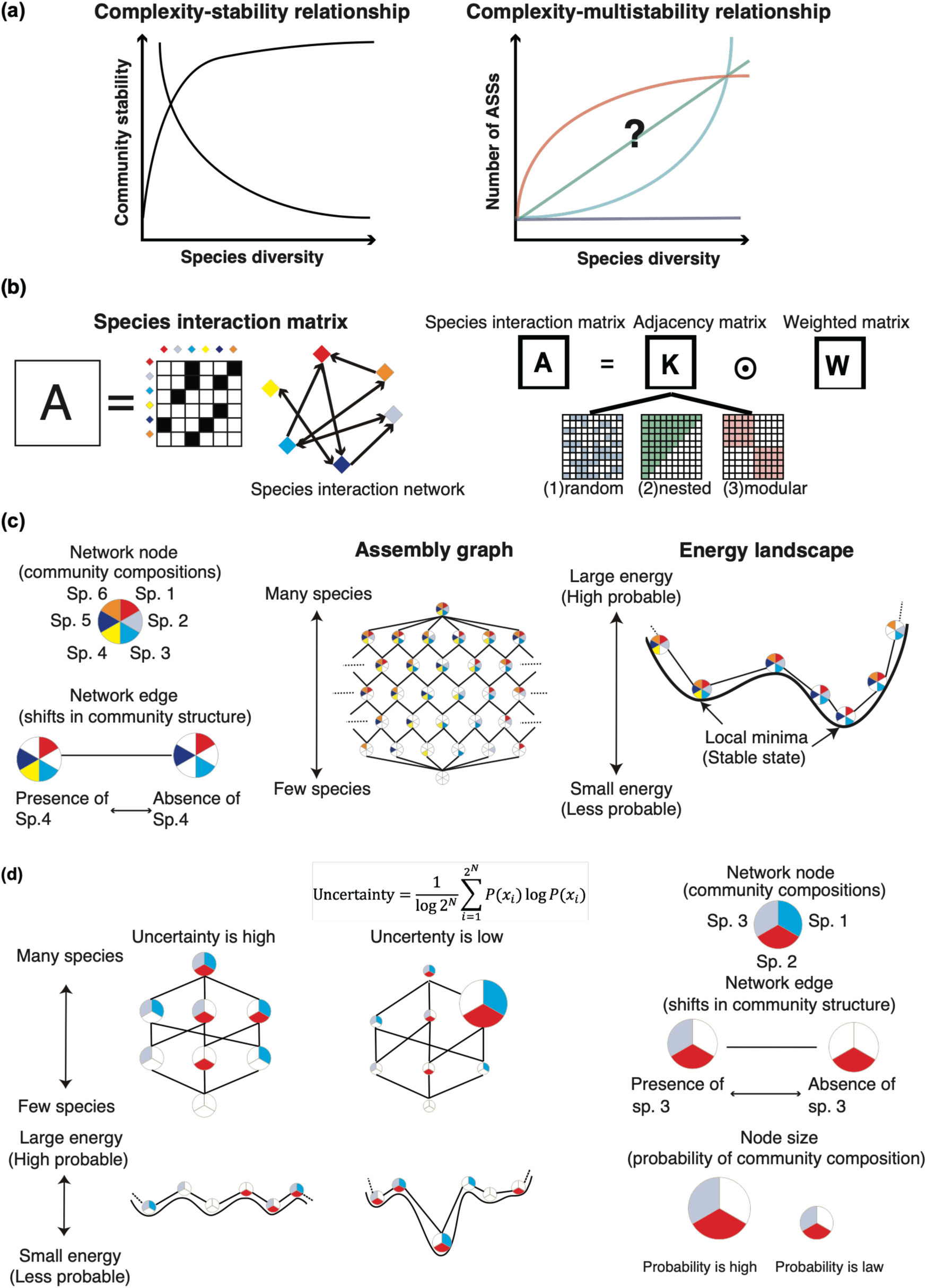
Conceptual overview and analytical framework. (a) Relationships between species diversity and community stability (left; the complexity–stability relationship) and between species diversity and the number of alternative stable states (ASSs) (right; the complexity-multistability relationship). (b) Species interaction networks are represented by interaction matrices, constructed as the Hadamard (element-wise) product of an adjacency matrix and a weighted matrix (Eq. 7). Three network topologies were considered for the adjacency matrix: random, nested, and modular. (c) Workflow for estimating the number of alternative stable states. The species interaction matrix is used to construct a stochastic community assembly model, from which the stationary distribution is obtained. The stationary distribution is then transformed into a stability landscape and combined with the community assembly graph to identify local minima, which correspond to alternative stable states. (d) Conceptual illustration of community uncertainty. Uncertainty is low when one community state dominates the stationary distribution and high when many community states have similar stationary probabilities. In the corresponding stability landscape, low uncertainty is associated with a single deep valley, whereas high uncertainty is characterized by many shallow valleys.

To address these limitations, we developed a stochastic community assembly framework that integrates stability landscape analysis, an approach inspired by energy landscape analysis (ELA; Suzuki et al. 2021, Toju et al. 2026). Previous studies typically relied on deterministic ordinary differential equation (ODE) models to quantify alternative stable states (e.g., Law and Morton 1993, Aguadé-Gorgorió et al. 2024), but these approaches require simulations from many initial conditions to identify all attractors, making them computationally demanding for species-rich communities (Song et al. 2021). In contrast, stochastic simulations allow transitions among community compositions, enabling estimation of the stationary distribution of community states without specifying numerous initial conditions. Stable states are identified as local minima in a stability landscape reconstructed from the stationary distribution of community states, following the principles of ELA (Ezaki et al. 2017, Suzuki et al. 2021, Masuda et al. 2025). ELA has recently been applied to empirical biological communities (Fujita et al. 2023, Kadoya et al. 2025, Noguchi et al. 2025, Hayashi et al. 2026), demonstrating the utility of stability landscape approaches in ecology. To make the framework computationally tractable, we focused on compositional multistability, defined by species presence–absence configurations rather than species abundances (Toju et al. 2026). We also quantified community uncertainty as the Shannon entropy of the stationary distribution over community states, which measures the long-term unpredictability of community assembly dynamics. This framework enables systematic quantification of multistability and community uncertainty across ecological communities of varying complexity.

In this study, we integrate a stochastic community assembly model with stability landscape analysis to investigate how community complexity influences multistability and community uncertainty. We apply this framework to a stochastic continent–island assembly model, in which colonization–extinction dynamics driven by species interactions generate transitions among community states. Stable states are identified as local minima in the resulting stability landscape. We examine how species interaction properties shape the relationship between species pool size and multistability, and whether multistability influences the uncertainty of long-term community composition. We show that mean interaction strength plays a key role in shaping the relationship between species pool size and multistability: when mean interaction strength is negative (i.e., under competitive interactions), larger species pools are associated with more alternative stable states. We further show that a greater number of alternative stable states does not necessarily increase community uncertainty, highlighting that uncertainty depends on the overall structure of the stability landscape rather than simply the number of stable states.

## 2 Methods

### 2.1 Model

We use a stochastic community assembly model with a continent–island structure, in which colonization from a regional species pool (the “continent”) and extinction within local communities (the “islands”) generate stochastic transitions among species presence–absence states. In classical island biogeography theory (MacArthur and Wilson 1967), colonization and extinction dynamics are primarily determined by island characteristics, whereas recent extensions incorporate species interactions into these processes, allowing community assembly to depend on biotic interactions (Gravel et al. 2011, Shibasaki and Terui 2024).

We consider a local community embedded within a regional species pool containing *N* species and characterized by an interspecific interaction matrix **A** = {*a_i_*_j_} (Fig. 1b). Let *C_i_*(*t*) denote the binary presence–absence of species *i* in the focal community at time *t*, where *C_i_*(*t*) = 1 indicates presence and *C_i_*(*t*) = 0 indicates absence. The colonization–extinction process is defined as:

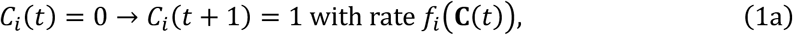

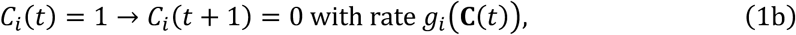

where *f_i_* and *g_i_* are the colonization and extinction rates, respectively. At each time step, a species is randomly selected, and its state is updated according to the corresponding transition rate. Both rates are functions of the current community state, **C**(*t*) = (*C*_1_(*t*), …, *C_N_*(*t*)).

Specifically, the colonization rate of species *i*, *f_i_*(**C**(*t*)), is modeled as a saturating function of the total strength of positive interspecific interactions with currently present species:

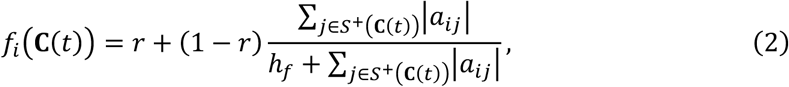

where *r* is the baseline colonization rate in the absence of positive interspecific interactions, *h_f_* is the half-saturation constant, *a_ij_* represents the effect of species *j* on species *i*, and *S*^+^(**C**(*t*)) denotes the set of species currently present that positively affect species *i* (*a_ij_* > 0) through facilitation, mutualism, or trophic interactions. The maximum colonization rate is scaled to 1.

Similarly, the extinction rate of species *i*, *g_i_*(**C**(*t*)), is modeled as a saturating function of the total strength of negative interspecific interactions with currently present species:

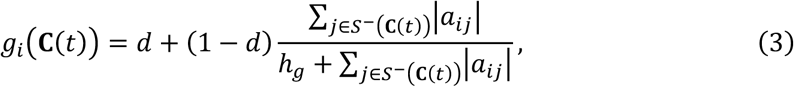

where *d* is the baseline extinction rate in the absence of negative interspecific interactions, *h_g_* is the half-saturation constant, *S*^-^(**C**(*t*)) denotes the set of currently present species that negatively affect species *i* (*a_ij_* < 0) through competition, predation, parasitism, or other antagonistic interactions. The maximum extinction rate is scaled to 1.

Together, these formulations provide a minimal representation of how biotic interactions shape species establishment and persistence during community assembly. Based on this model, we construct an assembly graph (Law and Morton 1993), where nodes represent community states and directed edges represent possible colonization or extinction transitions (Fig. 1c). The assembly graph therefore describes the complete state-transition structure of community dynamics.

### 2.2 Stability landscape analysis

The stochastic process defined by Eqs. 1–3 forms a Markov chain, in which transitions depend only on the current community state and not on previous states. When baseline colonization and extinction rates are positive (*r* > 0 and *d* > 0), species can colonize and become extinct independently of interspecific interactions. Under these conditions, the Markov chain converges to a unique stationary distribution (Appendix S1), which describes the long-term probability distribution of community states.

We define an energy-like index (*E*) for each community state **C** based on its stationary probability (*P*):

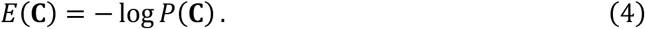

Because this transformation is inversely related to stationary probability, community states with high occurrence probabilities have low energy values, whereas rare states have high energy values. Following the ELA framework (Suzuki et al. 2021) and broader potential landscape framework (Dakos and Kéfi 2022), we interpret *E*(**C**) as the effective energy of a community state, with lower energy corresponding to higher stationary probability.

By combining the assembly graph with the energy values of each community state, we identify states whose energy is lower than that of all directly connected neighboring states in the assembly graph. These states correspond to local minima in the energy landscape, which are equivalent to local maxima in the stationary probability distribution. Following the potential landscape framework (Dakos and Kéfi 2022), these local minima represent valleys in the stability landscape and can be interpreted as stable community compositions. Therefore, the number of local minima provides an estimate of the number of alternative stable states (Fig. 1c). We implemented the local minima search using Rcpp.

### 2.3 Numerical simulations

Because the stochastic assembly process defines a Markov chain over all possible community states, the temporal dynamics of the probability distribution can be written as:

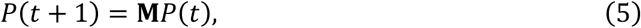

where *P*(t) is the probability distribution over community states at time *t*, and **M** is the transition matrix determined by Eqs. 1–3. The stationary distribution satisfies:

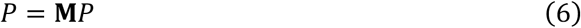

Thus, the stationary distribution can be obtained numerically as the normalized eigenvector of **M** corresponding to the eigenvalue 1. We then identified stable states (local minima) by combining the stationary distribution with the assembly graph following the stability landscape analysis described above (Eq. 4; Fig. 1b).

To investigate how species pool size (*N*) influences the number of alternative stable states under different interaction network topologies, we defined the *N* × *N* interaction matrix **A** as (Grilli et al. 2016):

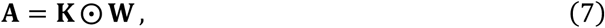

where **K** is an *N* × *N* adjacency matrix whose elements are either 0 or 1, **W** is an *N* × *N* weighted matrix, and ⊙ denotes the Hadamard (element-wise) product, such that *A_ij_* = *K_ij_W_ij_* (Fig. 1b). The matrix **K** specifies which pairs of species interact, whereas **W** determines the sign and strength of each interaction, where *A_ij_* represents the effect of species *j* on species *i*.

We assumed that **W** is a random matrix. Specifically, for each pair of interacting species *i* and *j*, the reciprocal interaction coefficients (*W_i_*_j_, *W*_j*i*_) were sampled independently from a bivariate normal distribution with means (*μ*, *μ*) and covariance matrix:

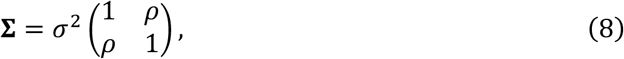

where *σ*^2^ is the variance and *ρ* is the correlation coefficient between reciprocal interaction coefficients. The parameter *μ* determines the mean interaction strength of the community, whereas *ρ* controls the symmetry of reciprocal interactions by determining the correlation between *W_ij_* and *W_ji_*.

In the numerical simulations, we considered three network topologies for the adjacency matrix **K**: (1) random, (2) fully nested, and (3) modular with equally sized modules (Fig. 1b). To isolate the effects of network topology, connectance was fixed at 0.5 across all networks. The mean *μ* was varied from −1 to 1 in increments of 0.25, the variance *σ*^2^ from 0.25 to 1 in increments of 0.25, and the correlation coefficient *ρ* from −1 to 1 in increments of 0.25. These parameter ranges encompass communities with predominantly negative interactions (competition), positive interactions (facilitation or cooperation), and asymmetric reciprocal interactions (predator–prey or parasite–host interactions). This resulted in 972 parameter combinations (9 means × 4 variances × 9 correlation coefficients × 3 network topologies). Species pool size (*N*) was varied from 5 to 20, and simulations were performed for all combinations of parameter values and species pool sizes. The remaining parameters were fixed at *r* = *d* = 0.1 and *h_f_* = *h_g_* = 1 (see Appendix S2 for additional parameter values). For each combination of parameters and species pool size, we generated 100 independent interaction networks and performed stochastic assembly simulations.

All numerical simulations were performed in R version 4.4.1 (R Core Team 2024) using the packages RSpectra version 0.16-2 (Qiu et al. 2024), Rcpp version 1.0.13 (Eddelbuettel et al. 2025), MASS version 7.3-61 (Venables and Ripley 2002), and Matrix version 1.7-0 (Bates et al. 2024). The stationary distribution was obtained as the eigenvector associated with the eigenvalue 1 of the stochastic transition matrix. Local minima were then identified from the normalized stationary distribution using the stability landscape analysis described above. To avoid numerical artifacts arising from finite numerical precision in the eigenvector calculation, only local minima with stationary probabilities greater than 10^−10^/∑*_i_ v_i_*, where *v* is the unnormalized eigenvector, were counted as alternative stable states. This threshold corresponds to the numerical precision of the eigenvector after normalization.

### 2.4 Uncertainty

To quantify uncertainty in community composition and examine its relationship with multistability, we used the normalized entropy of the stationary distribution (Song et al. 2021):

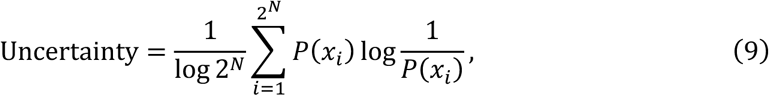

where *P*(*x_i_*) is the stationary probability of community state *x_i_*, representing a particular presence–absence configuration of the *N* species, **C** = (*C*, …, *C_N_*)), and 2*^N^*is the total number of possible community states. This metric corresponds to the Shannon entropy of the stationary distribution normalized by its maximum value, log 2*^N^* (Song 2025). It ranges from 0 to 1: values close to 0 indicate that community assembly is concentrated in a single state and therefore highly predictable, whereas values close to 1 indicate that many community states occur with similar probabilities and assembly outcomes are highly uncertain (Fig. 1d). Unlike simply counting alternative stable states, this metric incorporates the probability distribution across the entire stability landscape.

### 2.5 Relationship between species pool size and the number of alternative stable states

We calculated the number of alternative stable states for 972 combinations of species interaction parameters (9 means × 4 variances × 9 correlation coefficients × 3 network structures) across species pool sizes ranging from 5 to 20. For each parameter combination, we quantified the relationship between species pool size and the number of alternative stable states using Kendall’s rank correlation coefficient (*τ*), which captures monotonic relationships without assuming linearity. Parameter combinations with |*τ*| < 0.4 were classified as showing no clear monotonic relationship.

We applied a false discovery rate (FDR) correction across the 972 parameter combinations using a *q*-value threshold of 0.05. Relationships between species pool size and the number of alternative stable states were classified as positively monotonic when Kendall’s *τ* exceeded 0.4 and the *q*-value was < 0.05. For parameter combinations showing a significant monotonic relationship, we fitted three candidate models: a linear model, a generalized additive model (GAM) with a monotonic convex constraint, and a GAM with a monotonic concave constraint. The best-fitting model was selected using the Bayesian information criterion (BIC), and each relationship was classified as linear, convex, or concave.

To examine how interaction characteristics shape these relationship types, we used random forest classification (Breiman 2001) to predict whether the relationship between species pool size and the number of alternative stable states was linear, convex, concave, or other based on properties of the species interaction matrix, including the mean interaction strength, standard deviation, interaction symmetry, and network topology. We performed 100 random forest runs using inverse-frequency weighting to account for class imbalance. Classification performance and variable importance were averaged across runs. Variable importance was quantified as the mean decrease in accuracy, defined as the reduction in classification accuracy after randomly permuting the values of each predictor.

To assess the sensitivity of inferred relationship types to the maximum species pool size, we repeated the analysis after reducing the maximum species pool size from 20 to 18 or 15 species. We compared the resulting classifications of the relationship between species pool size and the number of alternative stable states with those obtained using the full range of species pool sizes (5-20 species). As in the main analysis, increasing relationships were identified using Kendall’s rank correlation coefficient and then classified as linear, convex, or concave based on BIC.

All statistical analyses were performed in R version 4.4.1 (R Core Team 2024) using the packages tidyverse version 2.0.0 (Wickham et al. 2019), scam version 1.2-18 (Pya 2025), and randomForest version 4.7-1.2 (Liaw and Wiener 2002).

### 2.6 Relationship between the number of alternative stable states and community uncertainty

We quantified the relationship between the number of alternative stable states and community uncertainty by calculating Kendall’s rank correlation coefficient (*τ*) for each of the 972 parameter combinations. We then examined how *τ* varied with mean interaction strength by plotting Kendall’s *τ* against the mean interaction strength. These analyses were performed in R version 4.4.1 (R Core Team 2024) using the package tidyverse version 2.0.0 (Wickham et al. 2019).

## 3 Results

### 3.1 Relationship between species pool size and the number of alternative stable states

We quantified the number of alternative stable states for 972 combinations of species interaction parameters across species pool sizes ranging from 5 to 20. The relationship between species pool size and the number of alternative stable states varied among parameter combinations, ranging from nearly constant to strongly increasing (Fig. 2a). Kendall’s rank correlation coefficient (*τ*) could not be computed for 15 parameter combinations because the number of alternative stable states did not vary across species pool sizes. For the remaining parameter combinations, Kendall’s *τ* ranged from −0.28 to 0.85 and exhibited a bimodal frequency distribution, with most parameter combinations exhibiting either weak negative or strong positive associations (Fig. 2a). Across all network topologies, mean interaction strength was negatively associated with Kendall’s *τ* (slope = −0.43 ± 0.01, 95% confidence intervals (CIs)) (Figs. 2b, S1–S9).

**Figure 2.**
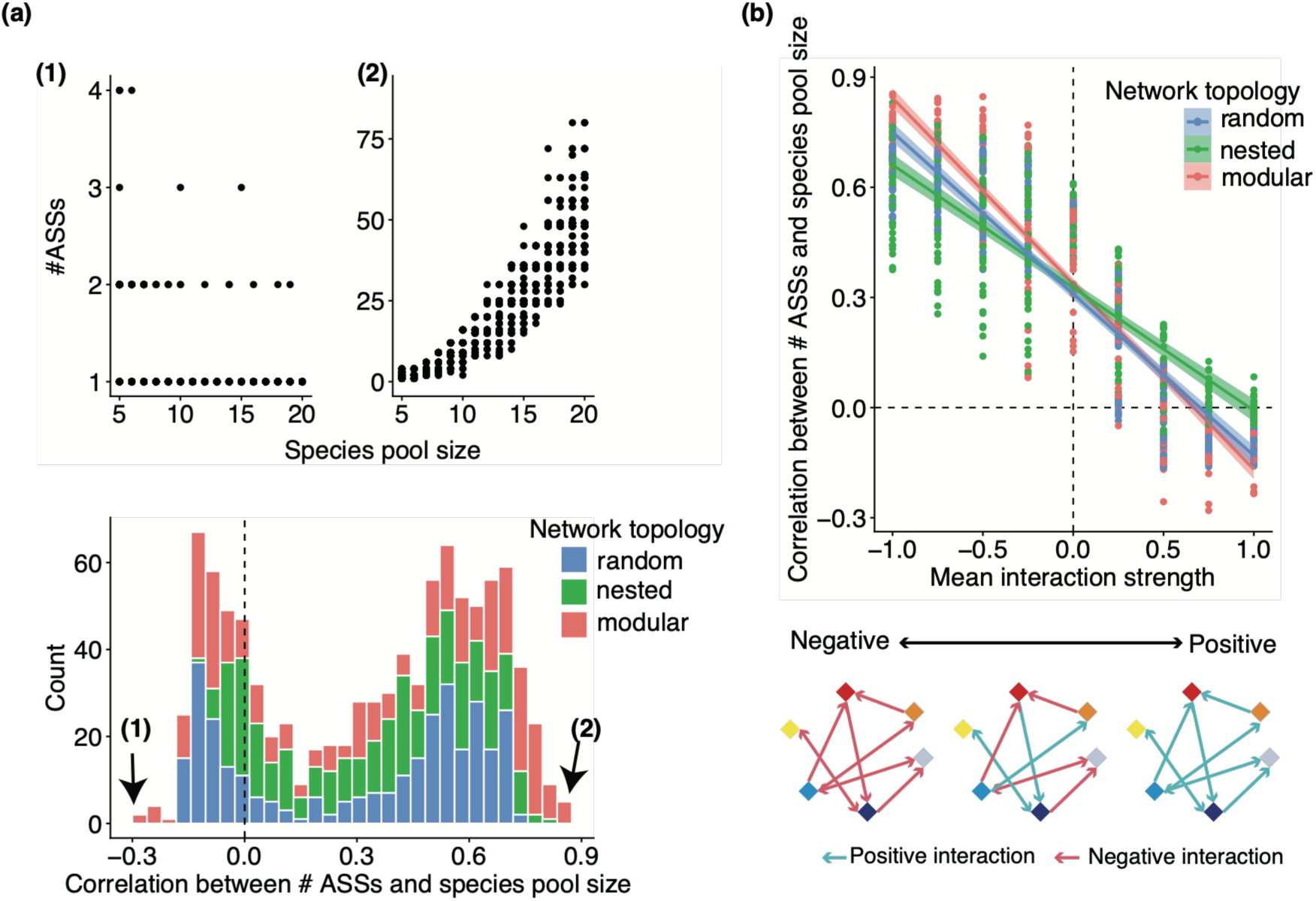
Relationship between species pool size and the number of alternative stable states across different interaction parameters. (a) The upper panels show representative examples of the relationship between species pool size and the number of alternative stable states (ASSs) for the parameter combinations yielding the smallest (a-1; *μ* = 0.75, *σ*^2^ = 0.5, *ρ* = 1, modular network) and largest (a-2; *μ* = −1, *σ*^2^ = 1, *ρ* = 1, modular network) values of Kendall’s rank correlation coefficient (*τ*). Here, *μ* denotes the mean interaction strength, *σ*^2^ denotes the variance of interaction strengths, and *ρ* denotes the correlation between reciprocal interaction strengths. The lower panel shows the distribution of Kendall’s *τ* across all parameter combinations. Colors indicate network topology (blue: random, green: nested, and red: modular). The dashed vertical line indicates *τ* = 0. (b) Relationship between mean interaction strength (*μ*) and Kendall’s *τ*. Solid lines show linear regressions and shaded areas indicate 95% confidence intervals (CIs). Colors denote network topology. The dashed horizontal line indicates *τ* = 0, and the dashed vertical line indicates *μ* = 0.

### 3.2 Increasing relationships between species pool size and the number of alternative stable states

Approximately half of the parameter combinations (484 of the 972) showed significant positive monotonic relationships between species pool size and the number of alternative stable states (Fig. 2a). Among these, BIC-based model selection classified 63 relationships as linear, 352 as convex, and 69 as concave. The remaining 488 parameter combinations showed either non-monotonic relationships or no significant relationship (Figs. 3a, S10a–S17a). Random forest classification achieved a mean accuracy of 82.36 ± 0.10% (95% CIs). Mean interaction strength was the most important predictor of relationship type (mean decrease in accuracy = 272.64 ± 2.15), followed by interaction symmetry (70.08 ± 0.53), network topology (61.54 ± 0.47), and the standard deviation of interaction strengths (13.05 ± 0.22) (Figs. 3c, S10b–S17b). Figure S18 shows how relationship classifications changed with the maximum species pool size. As the maximum species pool size increased, relationships were more frequently classified as convex rather than linear or concave (Figs. S19–S26).

**Figure 3.**
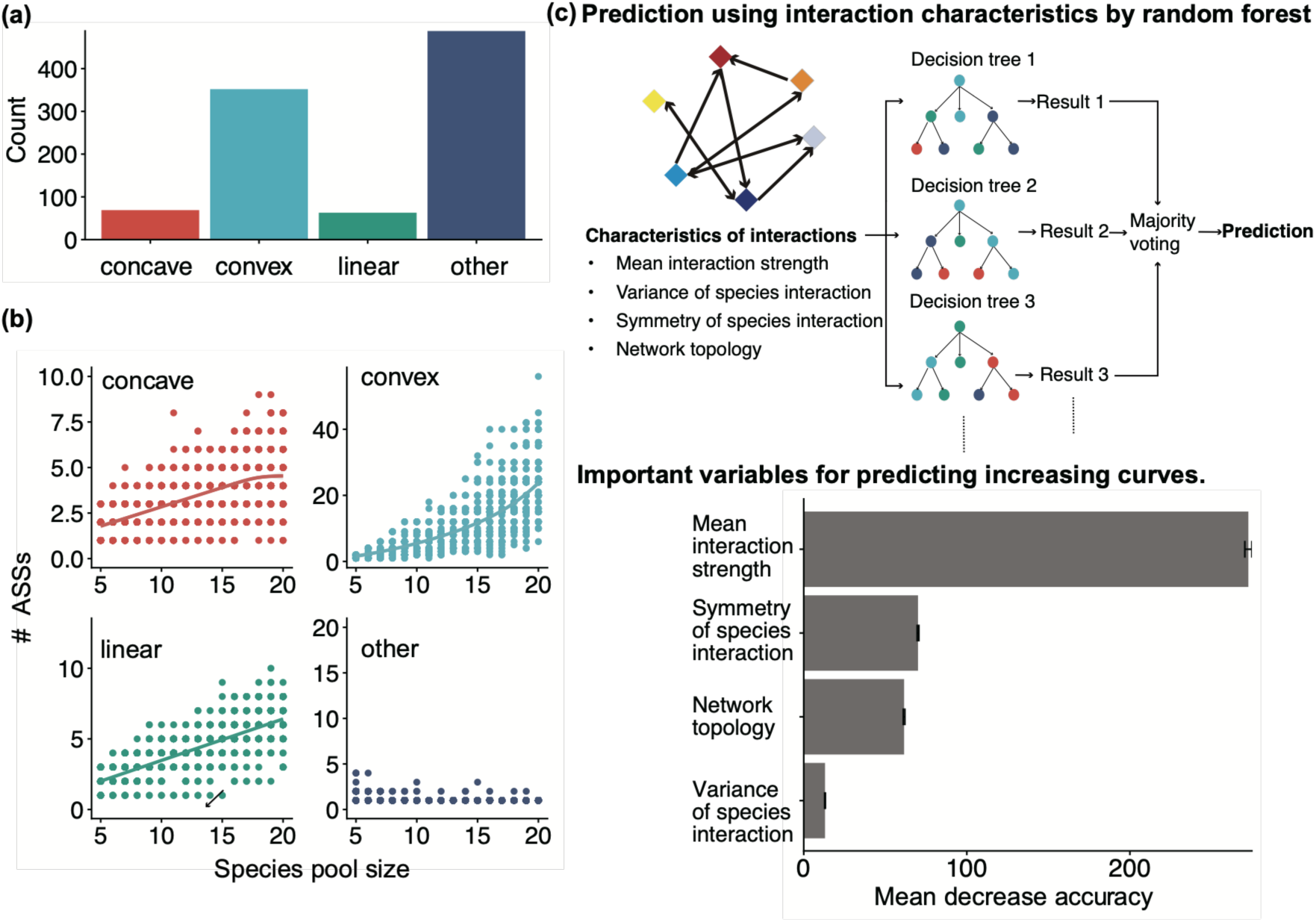
Classification of the relationships between species pool size and the number of alternative stable states. (a) Classification of relationship types for all 972 parameter combinations (red: concave, aqua: convex, green: linear, and dark blue: other). “Other” denotes parameter combinations that did not exhibit a significant monotonic increase. (b) Representative examples of the four relationship types: concave (*μ* = −0.75, *σ*^2^ = 0.75, *ρ* = 0, nested network), convex (*μ* = −0.5, *σ*^2^ = 1, *ρ* = 0.5, modular network), linear (*μ* = −1, *σ*^2^ = 0.25, *ρ* = −0.5, nested network), and other (*μ* = −1, *σ*^2^ = 1, *ρ* = 1, modular network). Here, *μ* denotes the mean interaction strength, *σ*^2^ denotes the variance of interaction strengths, and *ρ* denotes the correlation between reciprocal interaction strengths. Solid lines show fitted curves and shaded areas indicate 95% confidence intervals (CIs). (c) Upper panel: schematic of the random forest classifier used to classify relationship types from species interaction characteristics. Lower panel: variable importance measured as the mean decrease in accuracy after randomly permuting each predictor. Error bars indicate 95% CIs.

### 3.3 Relationship between the number of alternative stable states and community uncertainty

The relationship between the number of alternative stable states and community uncertainty (Eq. 9) varied among parameter combinations, ranging from positive to negative associations (Fig. 4a). Kendall’s *τ* ranged from −0.64 to 0.76 (Fig. 4a) and showed a quadratic relationship with mean interaction strength, reaching its minimum when mean interaction strength was close to −1 (Figs. 4b, S27–S35).

**Figure 4.**
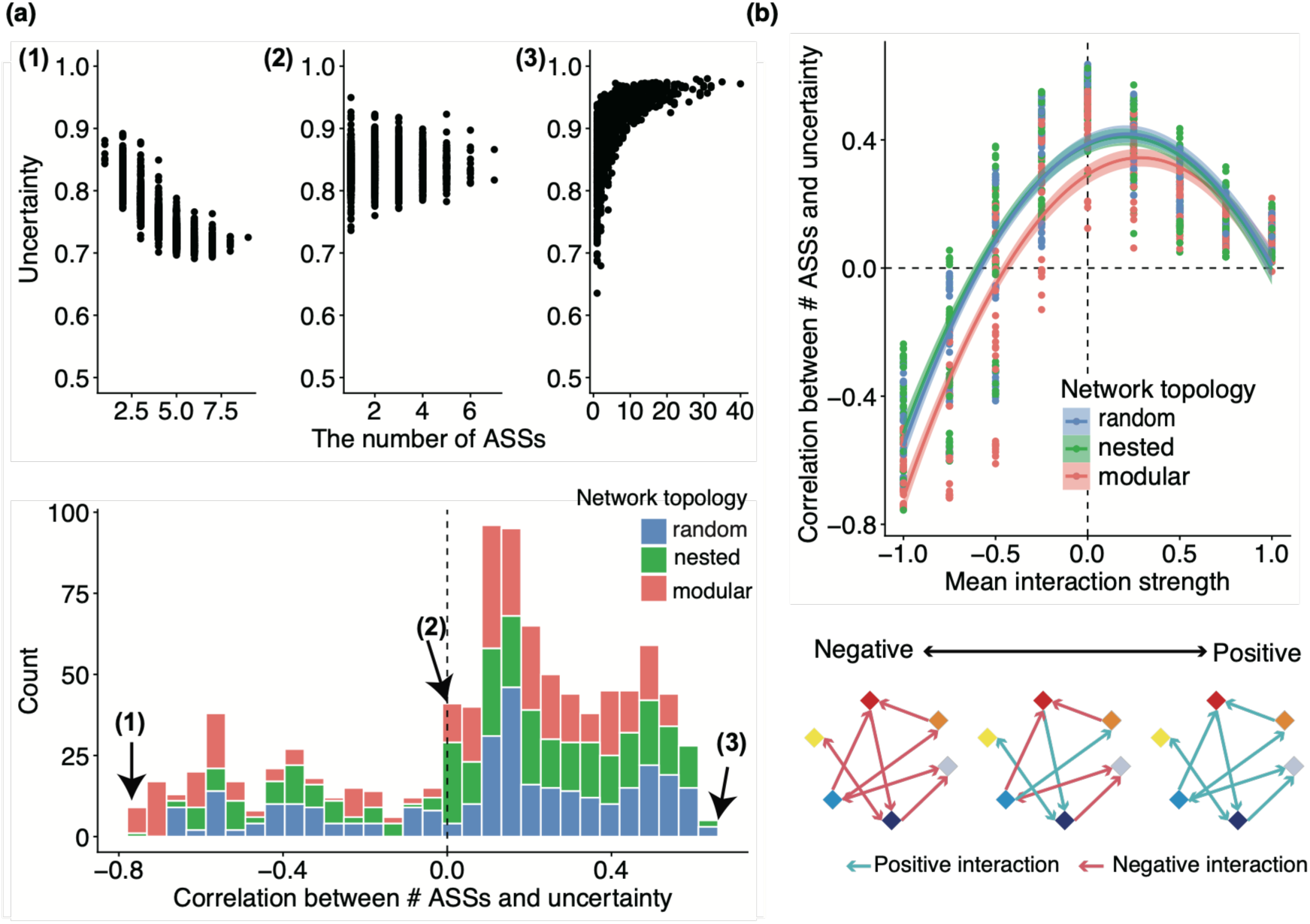
Relationship between the number of alternative stable states and community uncertainty across different interaction parameters. (a) The upper panels show representative examples of the relationship between the number of alternative stable states (ASSs) and community uncertainty for the parameter combinations yielding the smallest (a-1; *μ* = −1, *σ*^2^ = 0.25, *ρ* = 1, nested network), near-zero (a-2; *μ* = −0.5, *σ*^2^ = 0.5, *ρ* = 0.5, nested network), and largest (a-3; *μ* = 0, *σ*^2^ = 1, *ρ* = 0.75, random network) values of Kendall’s rank correlation coefficient (*τ*). Here, *μ* denotes the mean interaction strength, *σ*^2^ denotes the variance of interaction strengths, and *ρ* denotes the correlation between reciprocal interaction strengths. The lower panel shows the distribution of Kendall’s *τ* across all parameter combinations. Colors indicate network topology (blue: random, green: nested, and red: modular). The dashed vertical line indicates *τ* = 0. (b) Relationship between mean interaction strength (*μ*) and Kendall’s *τ*. Solid lines show quadratic regressions and shaded areas indicate 95% confidence intervals (CIs). Colors denote network topology. The dashed horizontal line indicates *τ* = 0, and the dashed vertical line indicates *μ* = 0.

## Discussion

Previous empirical studies have emphasized the importance of multistability in ecology (Scheffer et al. 2001, Beisner et al. 2003, Fukami 2015, Kéfi et al. 2019), whereas relatively few theoretical studies have explored multistability in species-rich communities (van Nes and Scheffer 2004, Kéfi et al. 2022, Aguadé-Gorgorió et al. 2024). This gap has partly reflected the computational cost of deterministic biomass-based models. To overcome this limitation, we developed a framework for quantifying multistability from species interaction matrices by integrating stochastic community assembly with stability landscape analysis (Fig. 1). Unlike conventional approaches focusing on local stability of individual equilibria (May 1972), our framework characterizes multistability as a system-level property by reconstructing the stability landscape of the entire community state space (Suzuki et al. 2021, Toju et al. 2026). By assigning an energy value to each community state according to its stationary probability, our approach extends landscape-based stability analysis to stochastic community assembly on discrete state spaces (Nolting and Abbott 2016). Our analyses revealed that species interaction properties strongly shape the relationship between species diversity and multistability, analogous to their well-established role in complexity–stability relationships (May 1972, Allesina and Tang 2012, Mougi and Kondoh 2012, Grilli et al. 2016, Giménez-Romero et al. 2025). In particular, although previous theoretical studies often suggested that the number of alternative stable states increases with species richness (Fried et al. 2016, Altieri et al. 2021, Aguadé-Gorgorió et al. 2024), 50.2% (488 of 972) of the parameter space did not show a monotonically increasing relationship (Fig. 2a). Moreover, our stochastic framework produced both convex and concave relationships between species pool size and the number of alternative stable states (Fig. 3a), demonstrating that these previously reported nonlinear patterns (Fried et al. 2016, Altieri et al. 2021) can be recovered within a unified stochastic framework.

Mean interaction strength emerged as the primary determinant of the relationship between species pool size and the number of alternative stable states within the parameter ranges considered here (Figs. 2b, S1). Under negative mean interaction strength, stronger competition amplified the positive relationship between species pool size and the number of alternative stable states. One possible explanation is that expanding the species pool increases the number of possible community configurations, whereas strong competition limits the number of species that can coexist locally. As a result, different subsets of the regional species pool can persist as alternative community configurations, increasing the number of stable states as the species pool expands. In our model, strong negative interactions increase extinction rates (Eq. 3), making community assembly more sensitive to species identity and arrival order. This is consistent with empirical evidence showing that competition amplifies priority effects (Vannette and Fukami 2014, Fukami 2015, Ke and Letten 2018).

In contrast, near-zero mean interaction strength weakened the relationship between species pool size and the number of alternative stable states compared with systems characterized by negative mean interaction strength (Fig. 2b). This pattern suggests that weakening competitive interactions may limit the increase in multistability associated with expanding species pools. This interpretation is broadly consistent with previous studies showing that trophic interactions can weaken priority effects by relaxing competitive exclusion among species (Morin 1984, Chase et al. 2009). When mean interaction strength was positive, the relationship between species pool size and the number of alternative stable states was close to zero. One possible explanation is that positive interactions can generate threshold-like, all-or-nothing dynamics, in which communities tend toward either coexistence or widespread exclusion, thereby limiting the emergence of additional alternative stable states as the species pool expands (Aguadé-Gorgorió et al. 2024). Such threshold dynamics have also been reported in mutualistic systems, including pollination networks that undergo abrupt collapse (Lever et al. 2014).

Species interaction properties also shaped the form of the relationship between species pool size and the number of alternative stable states. When mean interaction strength was non-positive, convex relationships were most frequently observed (Fig. 3), and their proportion increased as the maximum species pool size increased (Fig. S18). These results are consistent with previous studies reporting convex relationships between species richness and the number of stable states in communities characterized by a mixture of competitive and competitive– mutualistic interactions (Altieri et al. 2021, Aguadé-Gorgorió et al. 2024). In contrast, concave relationships were observed only under limited conditions in our analyses, even under negative mean interaction strengths (Figs. 3, S18). This discrepancy may reflect differences in model structure: studies reporting concave relationships assumed uniformly strong competition that inevitably led to competitive exclusion (Fried et al. 2016), whereas our model allowed interaction strengths to vary among species pairs. Thus, the shape of the diversity–multistability relationship depends not only on the mean interaction strength but also on how interaction strengths are distributed among species pairs.

We also found that a greater number of alternative stable states did not necessarily increase community uncertainty, a measure of the long-term unpredictability of community assembly dynamics (Song et al. 2021, Song 2025). Contrary to the intuitive expectation that community uncertainty increases with the number of alternative stable states, the relationship between the number of alternative stable states and community uncertainty ranged from negative to positive depending on the interaction parameters (Fig. 4a). This finding suggests that community uncertainty depends not only on the number of alternative stable states but also on their distribution across the stability landscape (Fig. 1d). Mean interaction strength was again the primary determinant of this relationship (Figs. 4b, S2). Kendall’s *τ* exhibited an approximately quadratic relationship with mean interaction strength, reaching a maximum near zero. This pattern suggests that when mean interaction strength is close to zero, multiple stable states may have similar stationary probabilities, resulting in a relatively flat stability landscape and high community uncertainty. As mean interaction strength became increasingly negative, a greater number of alternative stable states was associated with lower community uncertainty.

One possible explanation is that strong competition concentrates stationary probability on a limited number of community configurations while making alternative configurations increasingly unlikely. Thus, a community can have many alternative stable states while remaining relatively predictable if most stationary probability is concentrated in a subset of those states. Conversely, even a modest number of stable states can be associated with high uncertainty when their stationary probabilities are similar. These results demonstrate that understanding the predictability of long-term community composition requires characterizing the entire stability landscape rather than simply counting alternative stable states.

Although mean interaction strength was the strongest predictor of the diversity– multistability relationship, interaction symmetry, network topology, and variation in interaction strengths also contributed to determining the form of this relationship (Fig. 3c). This result indicates that the effects of species pool size on multistability cannot be explained solely by mean interaction strength. This parallels the complexity–stability relationship, in which the effects of increasing community complexity depend on multiple properties of species interaction networks rather than on species diversity alone (Okuyama and Holland 2008, Grilli et al. 2016). These findings suggest that empirical predictions of ecological multistability should consider not only the average strength of species interactions but also the architecture of interaction networks.

Our framework extends previous assembly-graph approaches by introducing a stochastic representation of community assembly and a stability landscape defined by the stationary distribution. Previous assembly-graph methods typically considered deterministic community dynamics and identified stable states by simulating trajectories from different initial conditions and counting the resulting absorbing states (Law and Morton 1993, Capitán et al. 2009, Serván and Allesina 2021). In contrast, our framework explicitly represents stochastic transitions among community states and reconstructs a stability landscape from their stationary distribution, allowing stable states to be identified as local minima. In this sense, our approach provides a quasi-potential-like representation for discrete assembly graphs, analogous to potential-based descriptions of stability landscapes in continuous systems (Nolting and Abbott 2016). More broadly, methods for quantifying multistability generally follow two complementary strategies, which identify stable states by simulating dynamics from multiple initial conditions (e.g., Gilpin and Case 1976, Aguadé-Gorgorió et al. 2024), and landscape-based approaches, which infer stable states from the underlying state space structure (Nolting and Abbott 2016). Our framework extends the latter perspective to discrete stochastic community assembly, thereby complementing conventional trajectory-based methods for studying ecological multistability.

Our framework has several limitations that point to directions for future research. First, community states are represented solely by species presence and absence, without accounting for continuous variables such as biomass or relative abundance. This compositional representation makes the analysis computationally tractable for species-rich communities, but it cannot capture abundance-mediated feedbacks or changes in interaction strength associated with species densities. Extending the framework to stochastic abundance-based models could therefore provide a way to analyze multistability while explicitly accounting for biomass dynamics. Second, we assumed that species interactions were time-invariant, whereas interaction strengths can change over ecological timescales in response to environmental variation (Poisot et al. 2012) or rapid evolution (Ellner 2013). Incorporating temporally varying interaction matrices would allow future studies to investigate how environmental and evolutionary processes reshape stability landscapes and alter the emergence, persistence, and diversity of alternative stable states. Third, our model represents the simplest case in which species interactions modify colonization and extinction rates, and does not explicitly distinguish among different ecological mechanisms underlying positive and negative interactions. Future extensions incorporating more mechanistic descriptions of species interactions, dispersal, and demographic processes could help determine how specific ecological mechanisms shape the structure of stability landscapes.

In summary, integrating stochastic community assembly with stability landscape analysis provides a general approach for studying multistability in complex ecological communities. By representing community assembly as a Markov process and reconstructing stability landscapes from species interaction matrices, our framework enables the identification and quantification of alternative stable states across the entire community state space. We show that the relationship between species pool size and multistability depends strongly on species interaction properties, with competitive interactions promoting the emergence of alternative community configurations as the species pool expands. Importantly, the number of alternative stable states alone does not determine community uncertainty: communities with many stable states can remain relatively predictable when stationary probability is concentrated on a subset of states, whereas communities with fewer stable states can be highly uncertain when probabilities are distributed more evenly. More broadly, our framework shifts the focus from simply counting alternative stable states to characterizing the structure of the stability landscape that governs long-term community composition. Because the framework requires only a species interaction matrix, it provides a potential basis for linking empirical interaction networks with multistability and community uncertainty, although reliable inference of interaction matrices remains an important challenge. Continued advances in ecological network inference may enable stability landscape analysis to be applied more broadly to empirical microbial, plant, and food-web communities, thereby enabling broader applications for linking ecological network structure to community assembly, multistability, and long-term predictability.

## Acknowledgements

We thank members of Yamamichi laboratory and Toju laboratory for their helpful comments on earlier version of the manuscript. We used ChatGPT for English proofreading.

