## Supplementary Materials for "Complexity-multistability relationships: How does species diversity shape the number of alternative stable states?"

Contents

Appendix S1. Existence and uniqueness of the stationary distribution of the continent–island model. 4

Appendix S2. Sensitivity analysis. 5

Figure S1. Relationships between species interaction properties and Kendall's rank correlation coefficient ($\tau$) between species pool size and the number of alternative stable states (ASSs). 7

Figure S2. Relationship between species pool size and the number of alternative stable states (ASSs) when $r = d = 0.1$ and $h_f = h_g = 0.1$. 8

Figure S3. Relationship between species pool size and the number of alternative stable states (ASSs) when $r = d = 0.1$ and $h_f = h_g = 10$. 9

Figure S4. Relationship between species pool size and the number of alternative stable states (ASSs) when $r = d = 0.3$ and $h_f = h_g = 0.1$. 10

Figure S5. Relationship between species pool size and the number of alternative stable states (ASSs) when $r = d = 0.3$ and $h_f = h_g = 1$. 11

Figure S6. Relationship between species pool size and the number of alternative stable states (ASSs) when $r = d = 0.3$ and $h_f = h_g = 10$. 12

Figure S7. Relationship between species pool size and the number of alternative stable states (ASSs) when $r = d = 0.6$ and $h_f = h_g = 0.1$. 13

Figure S8. Relationship between species pool size and the number of alternative stable states (ASSs) when $r = d = 0.6$ and $h_f = h_g = 1$. 14

Figure S9. Relationship between species pool size and the number of alternative stable states (ASSs) when $r = d = 0.6$ and $h_f = h_g = 10$. 15

Figure S10. Classification of increasing curves relationships between species pool size and the number of alternative stable states (ASSs) when $r = d = 0.1$ and $h_f = h_g = 0.1$. 16

Figure S11. Classification of increasing curves relationships between species pool size and the number of alternative stable states (ASSs) when $r = d = 0.1$ and $h_f = h_g = 10$. 17

Figure S12. Classification of increasing curves relationships between species pool size and the number of alternative stable states (ASSs) when $r = d = 0.3$ and $h_f = h_g = 0.1$. 18

Figure S13. Classification of increasing curves relationships between species pool size and the number of alternative stable states (ASSs) when $r = d = 0.3$ and $h_f = h_g = 1$. 19

Figure S14. Classification of increasing curves relationships between species pool size and the number of alternative stable states (ASSs) when $r = d = 0.3$ and $h_f = h_g = 10$. 20

Figure S15. Classification of increasing curves relationships between species pool size and the number of alternative stable states (ASSs) when $r = d = 0.6$ and $h_f = h_g = 0.1$. 21

Figure S16. Classification of increasing curves relationships between species pool size and the number of alternative stable states (ASSs) when $r = d = 0.6$ and $h_f = h_g = 1$. 22

Figure S17. Classification of increasing curves relationships between species pool size and the number of alternative stable states (ASSs) when $r = d = 0.6$ and $h_f = h_g = 10$. 23

Figure S18. Changes in relationship-type classifications with maximum species pool size. 24

Figure S19. Changes in relationship-type classifications with maximum species pool size when $r = d = 0.1$ and $h_f = h_g = 0.1$. 25

Figure S20. Changes in relationship-type classifications with maximum species pool size when $r = d = 0.1$ and $h_f = h_g = 10$. 26

Figure S21. Changes in relationship-type classifications with maximum species pool size when $r = d = 0.3$ and $h_f = h_g = 0.1$. 27

Figure S22. Changes in relationship-type classifications with maximum species pool size when $r = d = 0.3$ and $h_f = h_g = 1$. 28

Figure S23. Changes in relationship-type classifications with maximum species pool size when $r = d = 0.3$ and $h_f = h_g = 10$. 29

Figure S24. Changes in relationship-type classifications with maximum species pool size when $r = d = 0.6$ and $h_f = h_g = 0.1$. 30

Figure S25. Changes in relationship-type classifications with maximum species pool size when $r = d = 0.6$ and $h_f = h_g = 1$. 31

Figure S26. Changes in relationship-type classifications with maximum species pool size when $r = d = 0.6$ and $h_f = h_g = 10$. 32

Figure S27. Relationships between species interaction properties and Kendall's rank correlation coefficient ($\tau$) between the number of alternative stable states (ASSs) and community uncertainty. 33
Figure S28. Relationship between the number of alternative stable states (ASSs) and community uncertainty when $r = d = 0.6$ and $h_f = h_g = 0.1$. 34
Figure S29. Relationship between the number of alternative stable states (ASSs) and community uncertainty when $r = d = 0.1$ and $h_f = h_g = 10$. 35
Figure S30. Relationship between the number of alternative stable states (ASSs) and community uncertainty when $r = d = 0.3$ and $h_f = h_g = 0.1$. 36
Figure S31. Relationship between the number of alternative stable states (ASSs) and community uncertainty when $r = d = 0.3$ and $h_f = h_g = 1$. 37
Figure S32. Relationship between the number of alternative stable states (ASSs) and community uncertainty when $r = d = 0.3$ and $h_f = h_g = 10$. 38
Figure S33. Relationship between the number of alternative stable states (ASSs) and community uncertainty when $r = d = 0.6$ and $h_f = h_g = 0.1$. 39
Figure S34. Relationship between the number of alternative stable states (ASSs) and community uncertainty when $r = d = 0.6$ and $h_f = h_g = 1$. 40
Figure S35. Relationship between the number of alternative stable states (ASSs) and community uncertainty when $r = d = 0.6$ and $h_f = h_g = 10$. 41

***Appendix S1. Existence and uniqueness of the stationary distribution of the continent–island model.***

When the species pool size is $N$, the stochastic process defines a Markov chain on an $N$-dimensional hypercube. In this model, the neighborhood of a given community state consists of all states that differ by the presence or absence of a single species.

Assuming that one species is selected uniformly at random at each time step and may either colonize or go extinct, the transition probability from the current community state ($\mathbf{C}$) to a neighboring community state ($\mathbf{C}_{neighbor}$), which differs in the $i$-th species ($C_i$), is given by Eqs. 1-3 in the main text as follows:

$$\mathrm{Prob}\left(\mathbf{C} \to \mathbf{C}_{neighbor}\right) = \frac{1}{N}[(1 - C_i)f(\mathbf{C}) + C_i g(\mathbf{C})]. \qquad (A1)$$

From Eq. A1, the probability of remaining in the current community state is:

$$\mathrm{Prob}(\mathbf{C} \to \mathbf{C}) = 1 - \sum_{k \in neighbor} \mathrm{Prob}\,(\mathbf{C} \to \mathbf{C}_k) = 1 - \frac{1}{N}\sum_{i=1}^{N}[(1 - C_i)f(\mathbf{C}) + C_i g(\mathbf{C})]\,. \quad (A2)$$

Since the parameters $r$ and $d$ satisfy $0 < r, d < 1$, every colonization and extinction event has positive probability. Consequently, any community state can be reached from any other through a finite sequence of single-species transitions, implying that the Markov chain is irreducible. Furthermore, Eq. A2 gives a positive self-transition probability for every state, implying that the chain is aperiodic. Therefore, the Markov chain admits to a unique stationary distribution.

### *Appendix S2. Sensitivity analysis*

In the main text and Figures S1, S18, and S27, we assumed that $r = d = 0.1$ and $h_f = h_g = 1$. We conducted sensitivity analyses to evaluate the robustness of our results to these parameter values. To reduce computational cost, we assumed $r = d$ over 0.1, 0.3, and 0.6 and $h_f = h_g$ over 0.1, 1, and 10. We then repeated all analyses described in the main text for the resulting parameter combinations, excluding those already analyzed in the main manuscript.

#### S2-1. Relationship between species pool size and the number of alternative stable states (ASSs)

We quantified the relationship between species pool size and the number of ASSs for each parameter condition (Figs. S2–S9). Across all parameter combinations, the results were qualitatively consistent with the main analysis (Fig. 2). Kendall's rank correlation coefficients ranged from weakly negative to strongly positive and remained negatively associated with mean interaction strength.

#### S2-2. Classification of increasing relationships.

We classified the increasing relationships between species pool size and the number of ASSs for each parameter condition (Figs. S10a-S17a). Across all parameter combinations, convex relationships remained the dominant increasing relationship type, although the relative proportions of concave and linear relationships varied among parameter settings. Random forest classification consistently achieved mean accuracies exceeding 80%, with mean interaction strength remaining the most important predictor of relationship type across all parameter combinations (Figs. S10b-S17b). When $h_f = h_g = 0.1$, the ranking of the remaining predictors differed slightly from that in the main analysis, whereas the overall conclusions were unchanged.

#### S2-3. Effect of maximum species pool size.

We compared relationship classifications obtained using maximum species pool sizes of 15 and 20 (Figs. S19-S26). Across parameter combinations, increasing the maximum species pool size increased the proportion of convex relationships, consistent with the main analysis.

**S2-4. Relationship between the number of ASSs and community uncertainty**

We quantified the relationship between the number of ASSs and community uncertainty for each parameter condition (Figs. S28–S35). Across all parameter combinations, Kendall's rank correlation coefficients ranged from negative to strongly positive, consistent with the main analysis (Fig. 4). The approximately quadratic relationship between mean interaction strength and Kendall's rank correlation coefficient was also consistently observed, although increasing $h_f = h_g$ tended to make this relationship more linear.

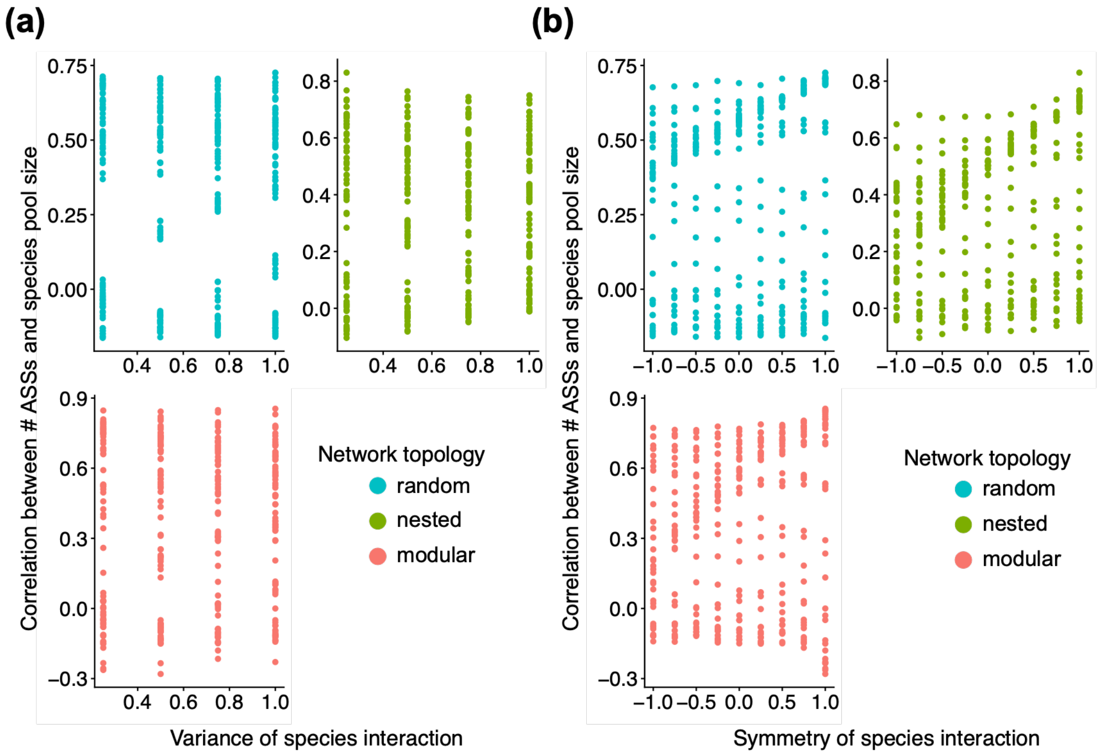


**Figure S1. Relationships between species interaction properties and Kendall's rank correlation coefficient ($\tau$) between species pool size and the number of alternative stable states (ASSs).** (a) Relationship between the variance of interaction strengths ($\sigma^2$) and Kendall's $\tau$. No clear association was observed (Pearson's correlation coefficient = 0.02). (b) Relationship between interaction symmetry ($\rho$) and Kendall's $\tau$. No clear association was observed (Pearson's correlation coefficient = 0.17). Colors indicate network topology (blue: random, green: nested, and red: modular).

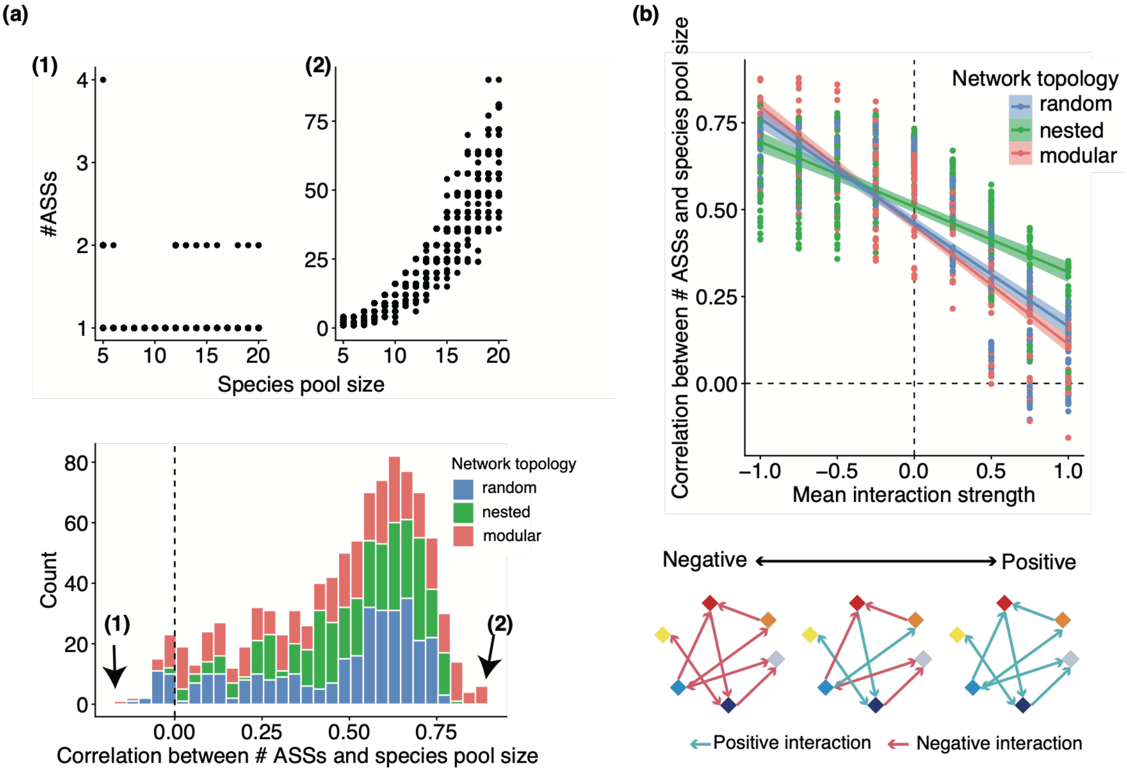


**Figure S2. Relationship between species pool size and the number of alternative stable states (ASSs) when $r = d = 0.1$ and $h_f = h_g = 0.1$.** (a) Kendall's rank correlation coefficient ($\tau$) ranged from −0.16 to 0.88. The upper panels show representative examples of the relationship between species pool size and the number of ASSs for the parameter combinations yielding the smallest (left; $\mu = 1, \sigma^2 = 0.25, \rho = 1$, modular network) and largest (right; $\mu = -0.75$, $\sigma^2 = 0.5, \rho = 1$, modular network) values of Kendall's $\tau$. The lower panel shows the distribution of Kendall's $\tau$ across all parameter combinations. Colors indicate network topology (blue: random, green: nested, and red: modular). The dashed vertical line indicates $\tau = 0$. (b) Relationship between mean interaction strength ($\mu$) and Kendall's $\tau$. Solid lines show linear regressions and shaded areas indicate 95% confidence intervals (CIs). Colors denote network topology. The dashed horizontal line indicates $\tau = 0$, and the dashed vertical line indicates $\mu = 0$. Pearson's correlation coefficients between Kendall's $\tau$ and interaction variance and symmetry were 0.16 and 0.18, respectively.

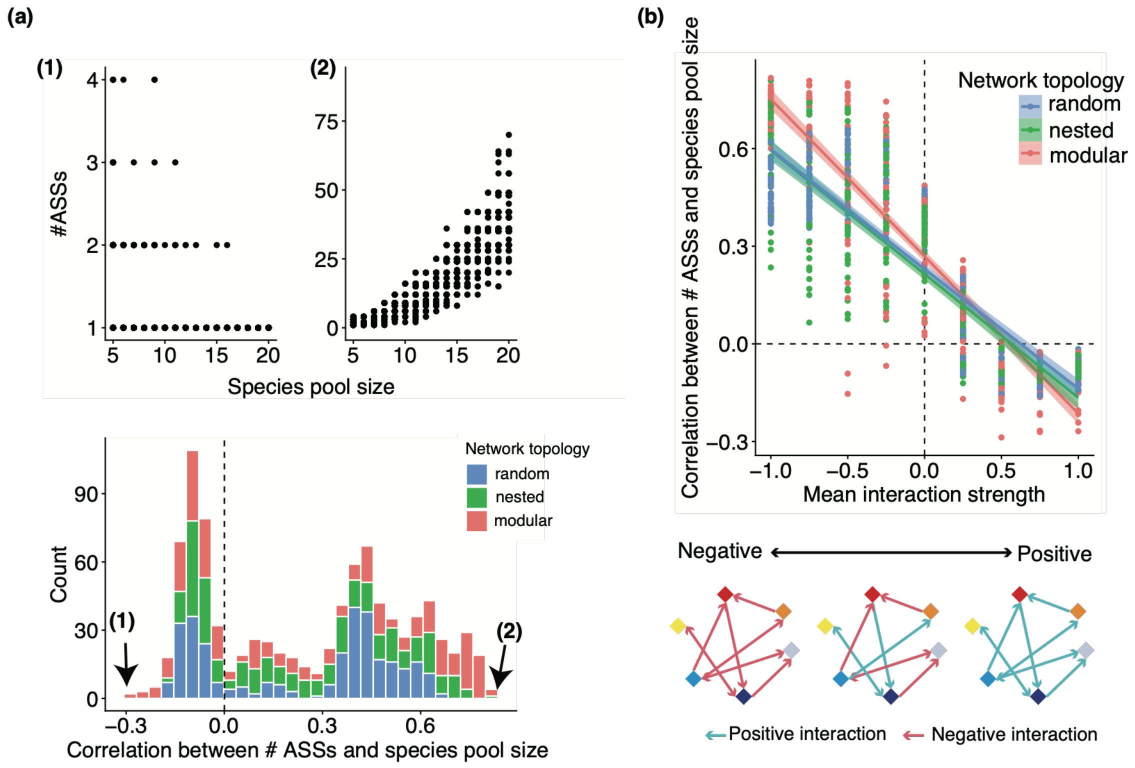


**Figure S3. Relationship between species pool size and the number of alternative stable states (ASSs) when $r = d = 0.1$ and $h_f = h_g = 10$.** (a) Kendall's rank correlation coefficient ($\tau$) ranged from −0.29 to 0.82. The upper panels show representative examples of the relationship between species pool size and the number of ASSs for the parameter combinations yielding the smallest (left; $\mu = 0.5, \sigma^2 = 0.25, \rho = 1$, modular network) and largest (right; $\mu = -1$, $\sigma^2 = 1, \rho = 1$, modular network) values of Kendall's $\tau$. The lower panel shows the distribution of Kendall's $\tau$ across all parameter combinations. Colors indicate network topology (blue: random, green: nested, and red: modular). The dashed vertical line indicates $\tau = 0$. (b) Relationship between mean interaction strength ($\mu$) and Kendall's $\tau$. Solid lines show linear regressions and shaded areas show 95% confidence intervals (CIs). Colors denote network topology. The dashed horizontal line indicates $\tau = 0$, and the dashed vertical line indicates $\mu = 0$. Pearson's correlation coefficients between Kendall's $\tau$ and interaction variance and symmetry were −0.06 and 0.21, respectively.

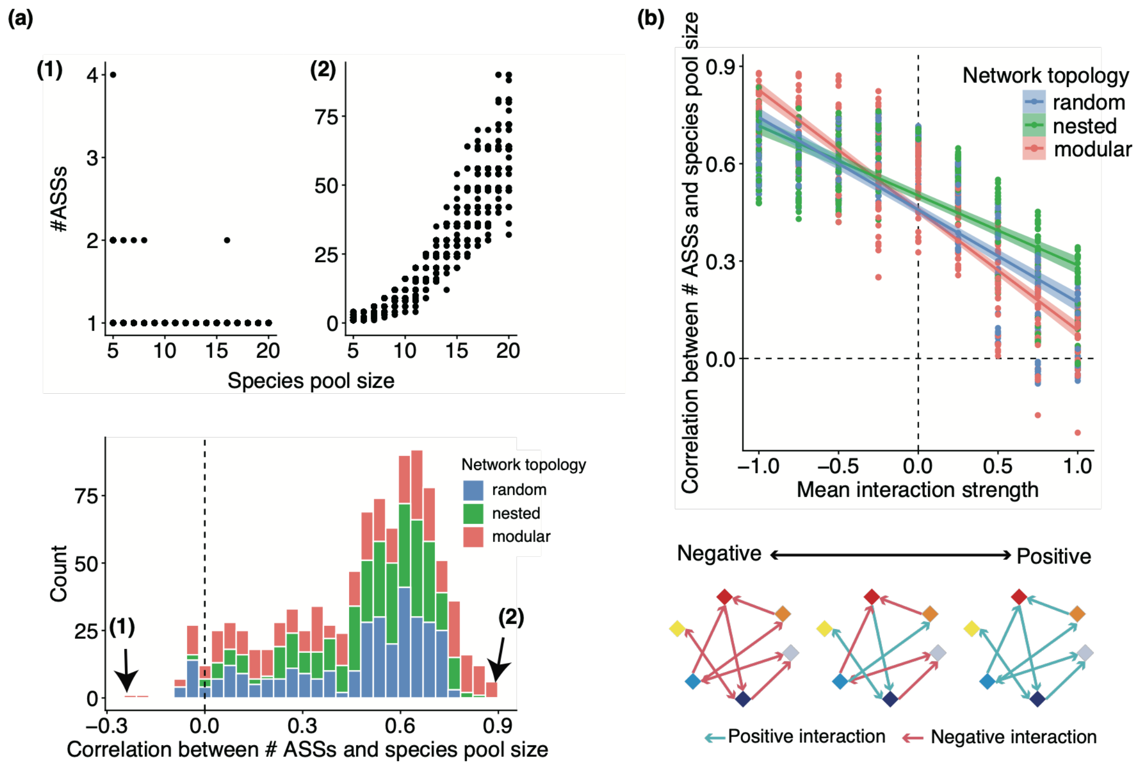


**Figure S4. Relationship between species pool size and the number of alternative stable states (ASSs) when $r = d = 0.3$ and $h_f = h_g = 0.1$.** (a) Kendall's rank correlation coefficient ($\tau$) ranged from −0.23 to 0.88. The upper panels show representative examples of the relationship between species pool size and the number of ASSs for the parameter combinations yielding the smallest (left; $\mu = 1, \sigma^2 = 0.25, \rho = 1$, modular network) and largest (right; $\mu = -1,\ \sigma^2 = 1, \rho = 1$, modular network) values of Kendall's $\tau$. The lower panel shows the distribution of Kendall's $\tau$ across all parameter combinations. Colors indicate network topology (blue: random, green: nested, and red: modular). The dashed vertical line indicates $\tau = 0$. (b) Relationship between mean interaction strength ($\mu$) and Kendall's $\tau$. Solid lines show linear regressions and shaded areas indicate 95% confidence intervals (CIs). Colors denote network topology. The dashed horizontal line indicates $\tau = 0$, and the dashed vertical line indicates $\mu = 0$. Pearson's correlation coefficients between Kendall's $\tau$ and interaction variance and symmetry were 0.15 and 0.15, respectively.

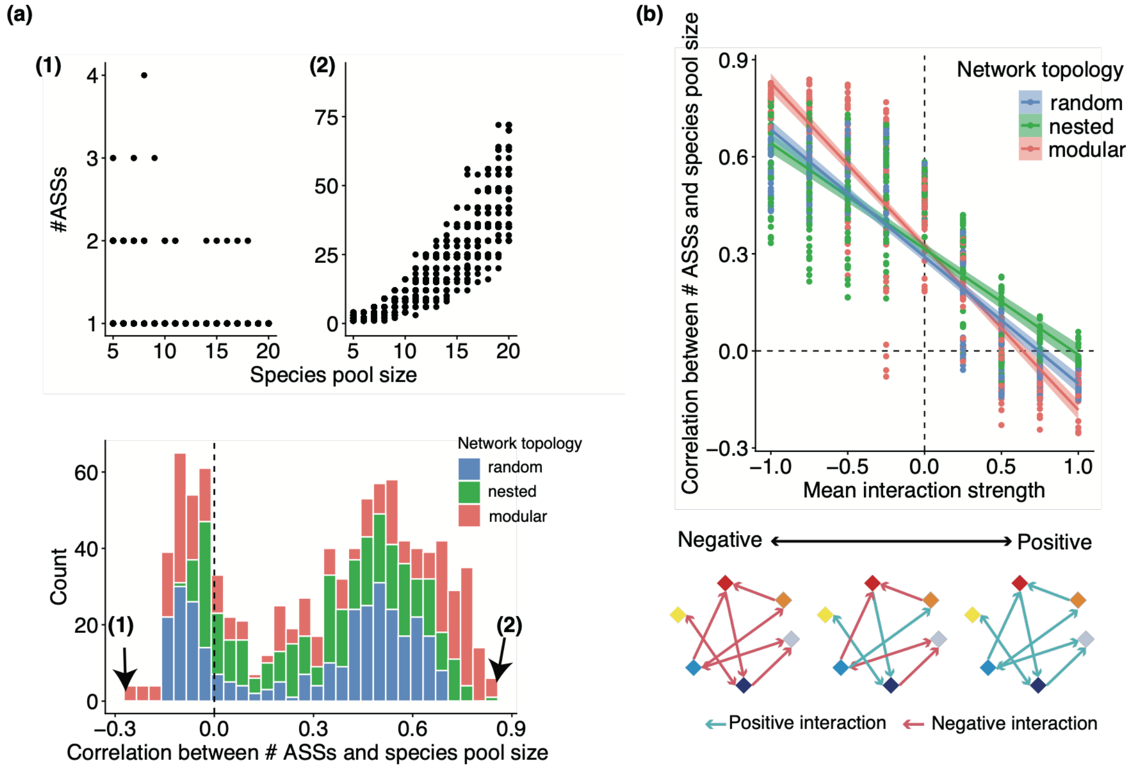


**Figure S5. Relationship between species pool size and the number of alternative stable states (ASSs) when $r = d = 0.3$ and $h_f = h_g = 1$.** (a) Kendall's rank correlation coefficient ($\tau$) ranged from −0.25 to 0.84. The upper panels show representative examples of the relationship between species pool size and the number of ASSs for the parameter combinations yielding the smallest (left; $\mu = 1, \sigma^2 = 0.75, \rho = 1$, modular network) and largest (right; $\mu = -0.75,\ \sigma^2 = 0.75, \rho = 1$, modular network) values of Kendall's $\tau$. The lower panel shows the distribution of Kendall's $\tau$ across all parameter combinations. Colors indicate network topology (blue: random, green: nested, and red: modular). The dashed vertical line indicates $\tau = 0$. (b) Relationship between mean interaction strength ($\mu$) and Kendall's $\tau$. Solid lines show linear regressions and shaded areas indicate 95% confidence intervals (CIs). Colors denote network topology. The dashed horizontal line indicates $\tau = 0$, and the dashed vertical line indicates $\mu = 0$. Pearson's correlation coefficients between Kendall's $\tau$ and interaction variance and symmetry were 0.003 and 0.18, respectively.

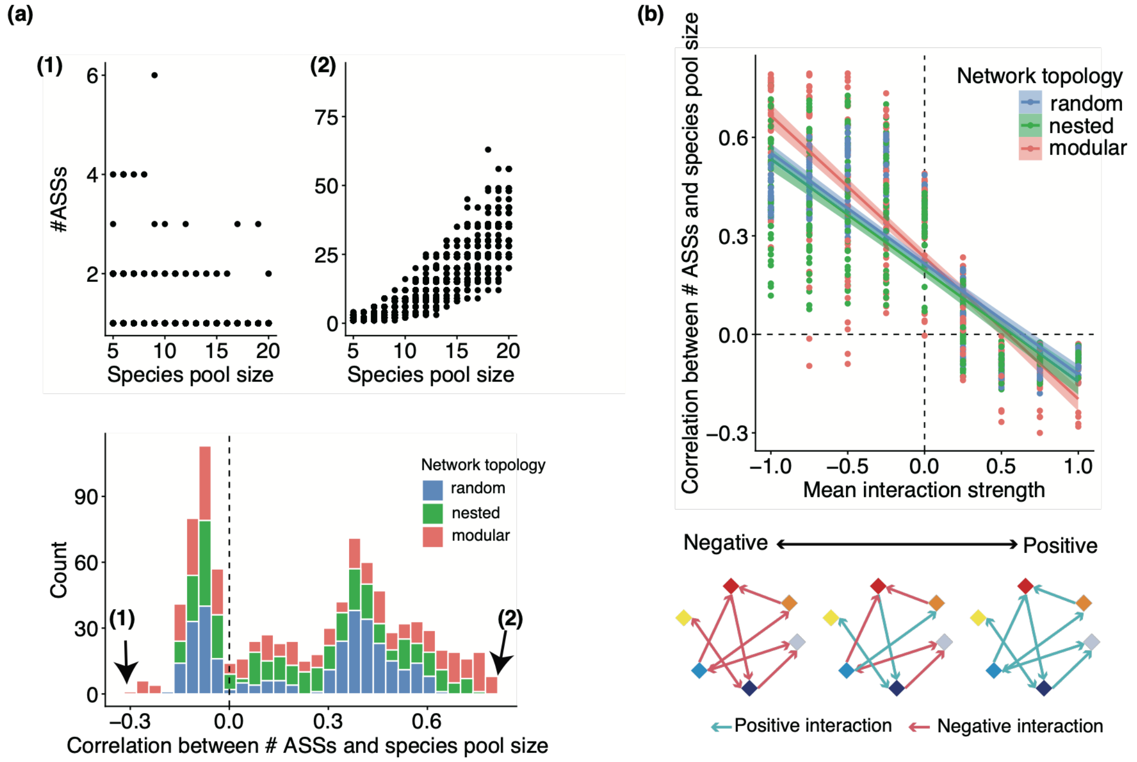


**Figure S6. Relationship between species pool size and the number of alternative stable states (ASSs) when $r = d = 0.3$ and $h_f = h_g = 10$.** (a) Kendall's rank correlation coefficient ($\tau$) ranged from −0.30 to 0.79. The upper panels show representative examples of the relationship between species pool size and the number of ASSs for the parameter combinations yielding the smallest (left; $\mu = 0.75, \sigma^2 = 0.75, \rho = 1$, modular network) and largest (right; $\mu = -0.5$, $\sigma^2 = 0.25, \rho = 1$, modular network) values of Kendall's $\tau$. The lower panel shows the distribution of Kendall's $\tau$ across all parameter combinations. Colors indicate network topology (blue: random, green: nested, and red: modular). The dashed vertical line indicates $\tau = 0$. (b) Relationship between mean interaction strength ($\mu$) and Kendall's $\tau$. Solid lines show linear regressions and shaded areas indicate 95% confidence intervals (CIs). Colors denote network topology. The dashed horizontal line indicates $\tau = 0$, and the dashed vertical line indicates $\mu = 0$. Pearson's correlation coefficients between Kendall's $\tau$ and interaction variance and symmetry were −0.06 and 0.23, respectively.

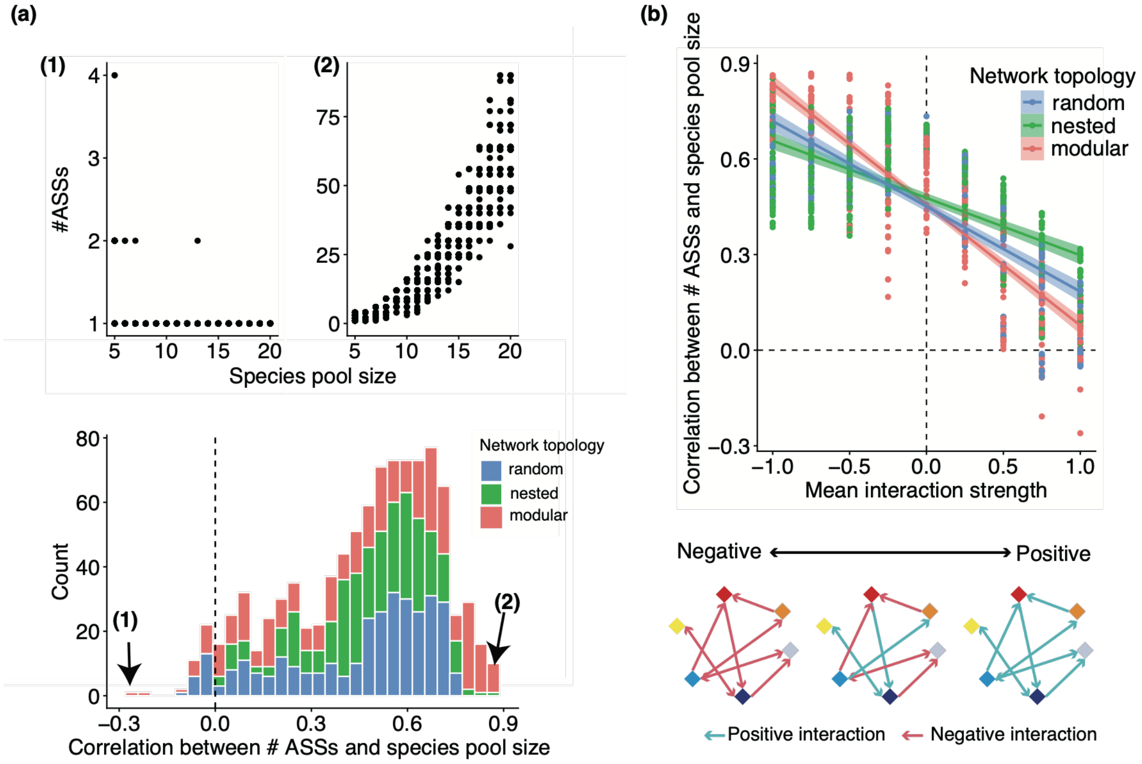


**Figure S7. Relationship between species pool size and the number of alternative stable states (ASSs) when $r = d = 0.6$ and $h_f = h_g = 0.1$.** (a) Kendall's rank correlation coefficient ($\tau$) ranged from −0.26 to 0.87. The upper panels show representative examples of the relationship between species pool size and the number of ASSs for the parameter combinations yielding the smallest (left; $\mu = 1, \sigma^2 = 0.25, \rho = 1$, modular network) and largest (right; $\mu = -0.75$, $\sigma^2 = 0.75, \rho = 1$, modular network) values of Kendall's $\tau$. The lower panel shows the distribution of Kendall's $\tau$ across all parameter combinations. Colors indicate network topology (blue: random, green: nested, and red: modular). The dashed vertical line indicates $\tau = 0$. (b) Relationship between mean interaction strength ($\mu$) and Kendall's $\tau$. Solid lines show linear regressions and shaded areas indicate 95% confidence intervals (CIs). Colors denote network topology. The dashed horizontal line indicates $\tau = 0$, and the dashed vertical line indicates $\mu = 0$. Pearson's correlation coefficients between Kendall's $\tau$ and interaction variance and symmetry were 0.14 and 0.16, respectively.

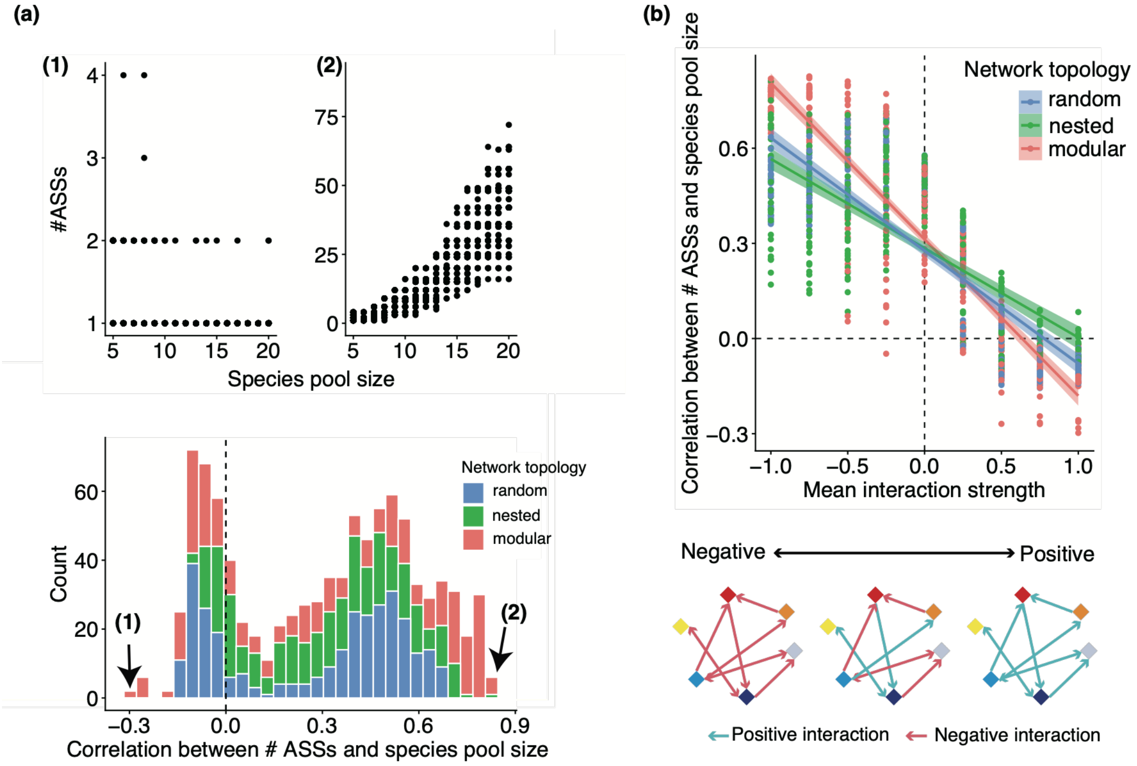


**Figure S8. Relationship between species pool size and the number of alternative stable states (ASSs) when $r = d = 0.6$ and $h_f = h_g = 1$.** (a) Kendall's rank correlation coefficient ($\tau$) ranged from −0.30 to 0.83. The upper panels show representative examples of the relationship between species pool size and the number of ASSs for the parameter combinations yielding the smallest (left; $\mu = 1, \sigma^2 = 0.75, \rho = 1$, modular network) and largest (right; $\mu = -0.75,\ \sigma^2 = 0.75, \rho = 1$, modular network) values of Kendall's $\tau$. The lower panel shows the distribution of Kendall's $\tau$ across all parameter combinations. Colors indicate network topology (blue: random, green: nested, and red: modular). The dashed vertical line indicates $\tau = 0$. (b) Relationship between mean interaction strength ($\mu$) and Kendall's $\tau$. Solid lines show linear regressions and shaded areas indicate 95% confidence intervals (CIs). Colors denote network topology. The dashed horizontal line indicates $\tau = 0$, and the dashed vertical line indicates $\mu = 0$. Pearson's correlation coefficients between Kendall's $\tau$ and interaction variance and symmetry were 0.02 and 0.20, respectively.

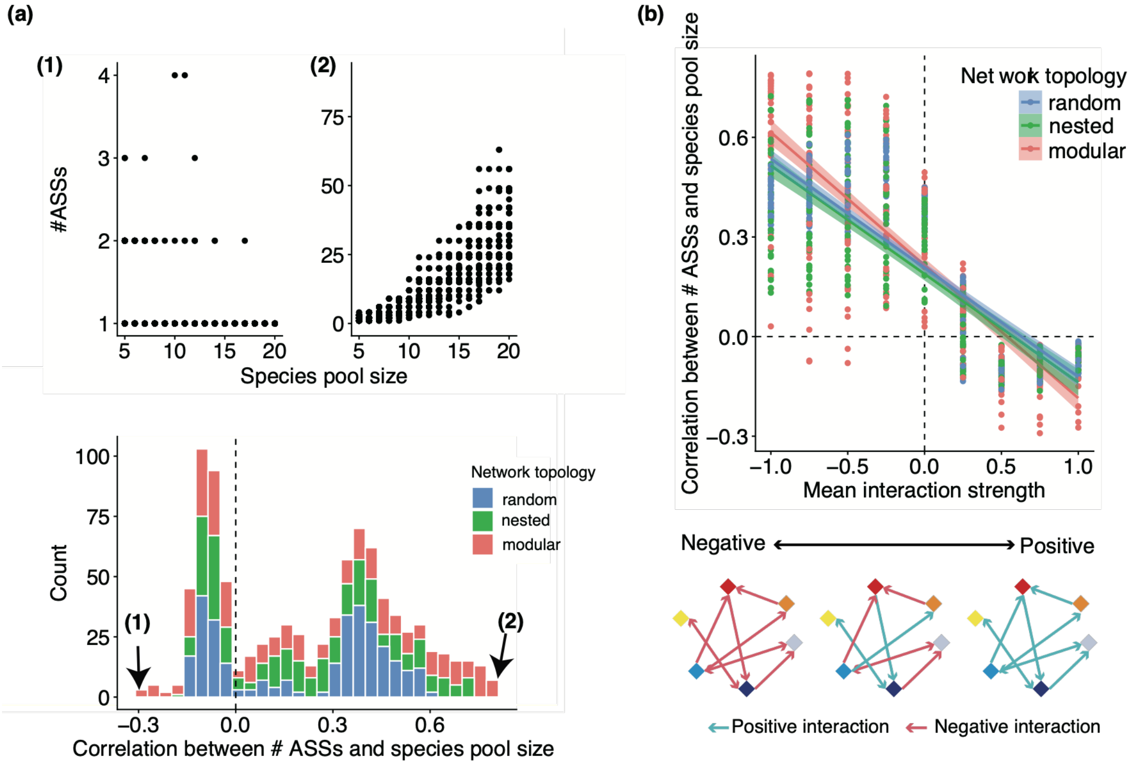


**Figure S9. Relationship between species pool size and the number of alternative stable states (ASSs) when $r = d = 0.6$ and $h_f = h_g = 10$.** (a) Kendall's rank correlation coefficient ($\tau$) ranged from −0.29 to 0.79. The upper panels show representative examples of the relationship between species pool size and the number of ASSs for the parameter combinations yielding the smallest (left; $\mu = 0.75, \sigma^2 = 0.5, \rho = 1$, modular network) and largest (right; $\mu = -0.75$, $\sigma^2 = 0.5, \rho = 1$, modular network) values of Kendall's $\tau$. The lower panel shows the distribution of Kendall's $\tau$ across all parameter combinations. Colors indicate network topology (blue: random, green: nested, and red: modular). The dashed vertical line indicates $\tau = 0$. (b) Relationship between mean interaction strength ($\mu$) and Kendall's $\tau$. Solid lines show linear regressions and shaded areas indicate 95% confidence intervals (CIs). Colors denote network topology. The dashed horizontal line indicates $\tau = 0$, and the dashed vertical line indicates $\mu = 0$. Pearson's correlation coefficients between Kendall's $\tau$ and interaction variance and symmetry were −0.05 and 0.24, respectively.

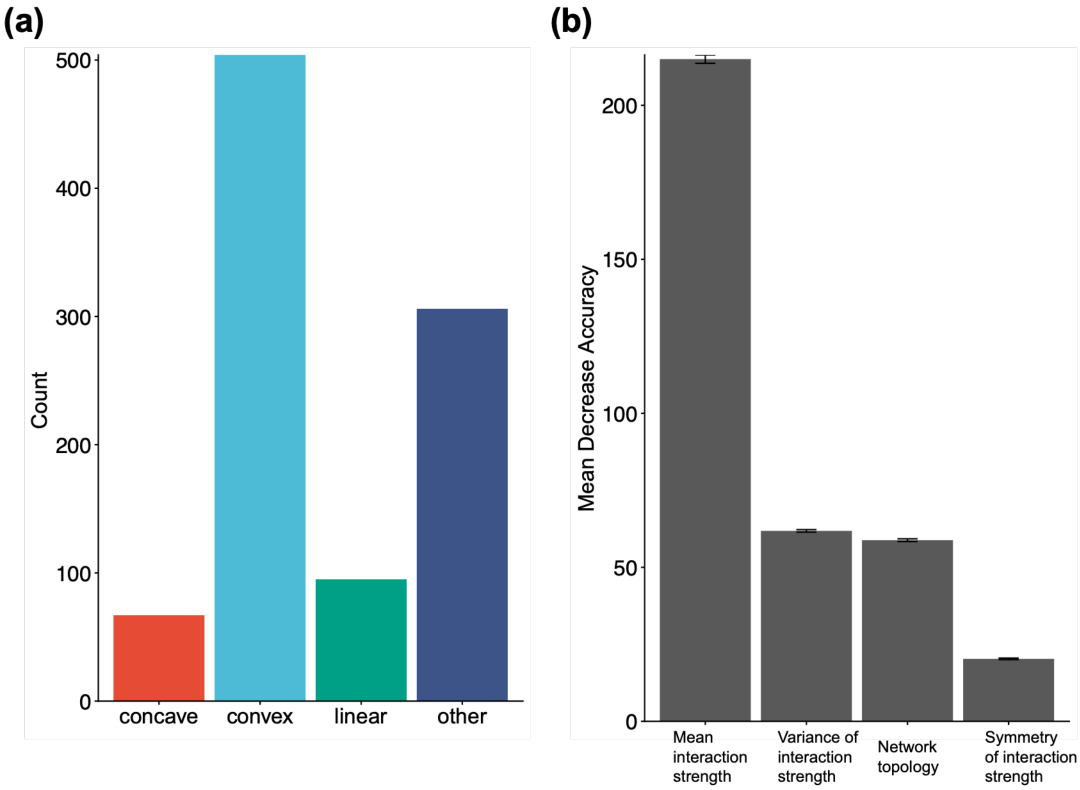


**Figure S10. Classification of increasing relationships between species pool size and the number of alternative stable states (ASSs) when $r = d = 0.1$ and $h_f = h_g = 0.1$.** The left panel shows the relationship-type classification for all 972 parameter combinations (concave: 95, convex: 504, linear: 95, and other: 306). The right panel shows the variable importance for classifying relationship types based on species interaction characteristics. Variable importance was quantified as the mean decrease in prediction accuracy after randomly permuting each predictor. The Y-axis represents the mean decrease in prediction accuracy. Error bars indicate 95% confidence intervals (CIs). The mean classification accuracy was 80.45 ± 0.10% (95% CIs).

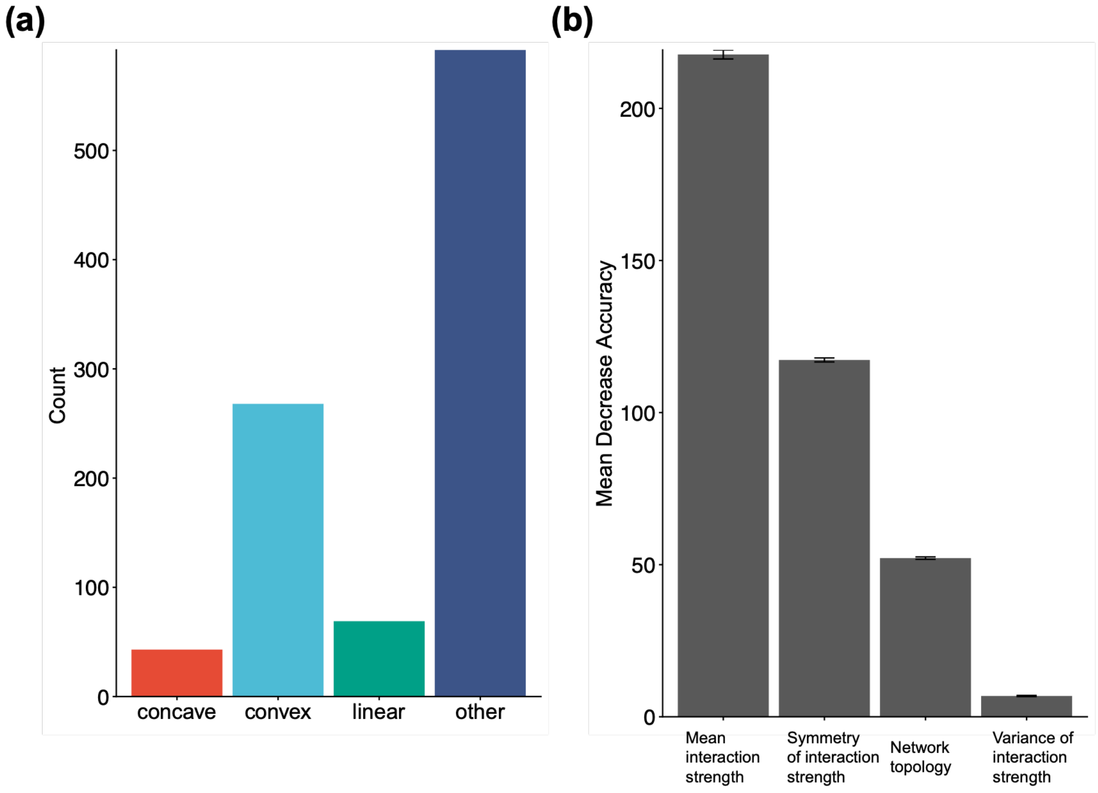


**Figure S11. Classification of increasing relationships between species pool size and the number of alternative stable states (ASSs) when $r = d = 0.1$ and $h_f = h_g = 10$.** The left panel shows the relationship-type classification for all 972 parameter combinations (concave: 43, convex: 268, linear: 69, and other: 592). The right panel shows the variable importance for classifying relationship types based on species interaction characteristics. Variable importance was quantified as the mean decrease in prediction accuracy after randomly permuting each predictor. The Y-axis represents the mean decrease in prediction accuracy. Error bars indicate 95% confidence intervals (CIs). The mean classification accuracy was 83.10 ± 0.10% (95% CIs).

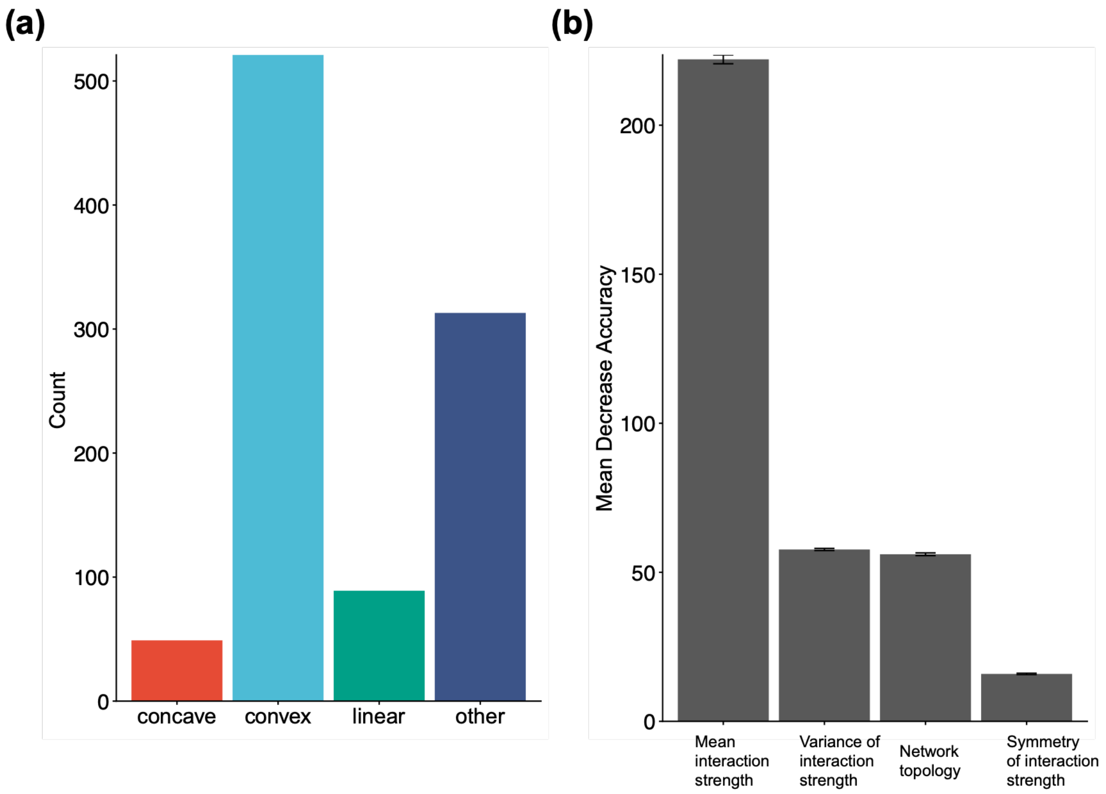


**Figure S12. Classification of increasing relationships between species pool size and the number of alternative stable states (ASSs) when $r = d = 0.3$ and $h_f = h_g = 0.1$.** The left panel shows the relationship-type classification for all 972 parameter combinations (concave: 49, convex: 521, linear: 89, and other: 313). The right panel shows the variable importance for classifying relationship types based on species interaction characteristics. Variable importance was quantified as the mean decrease in prediction accuracy after randomly permuting each predictor. The Y-axis represents the mean decrease in prediction accuracy. Error bars indicate 95% confidence intervals (CIs). The mean classification accuracy was 82.17 ± 0.09% (95% CIs).

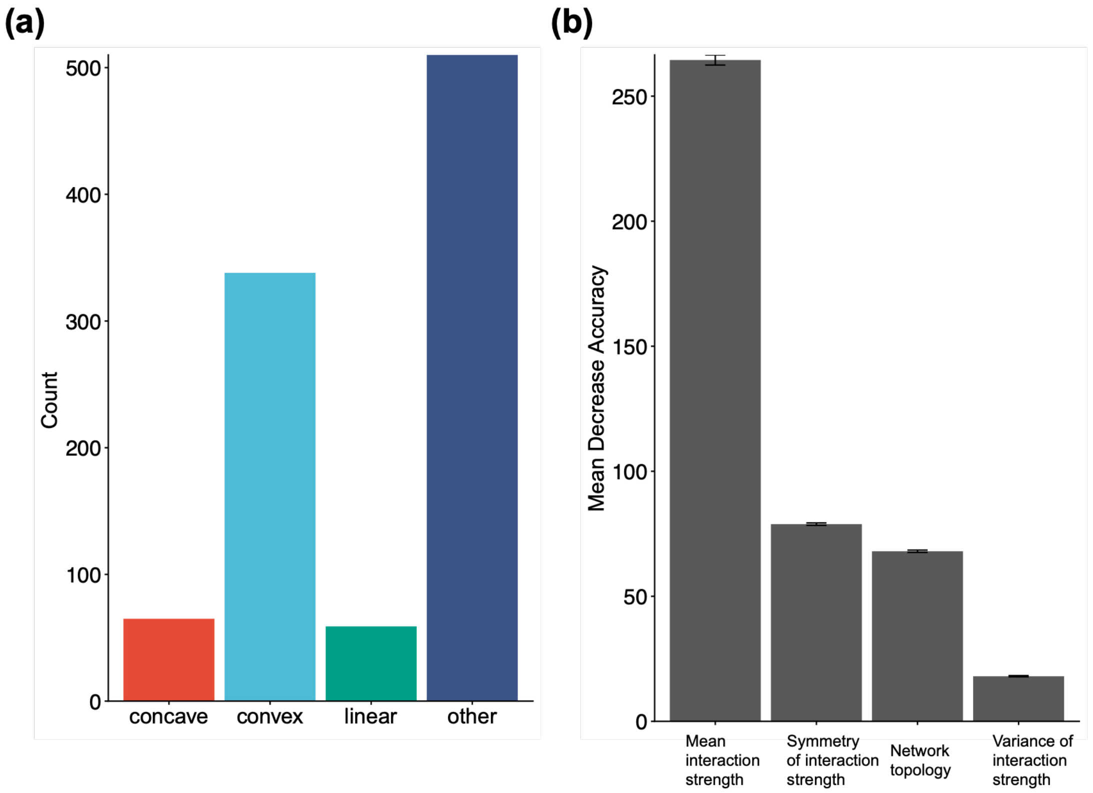


**Figure S13. Classification of increasing relationships between species pool size and the number of alternative stable states (ASSs) when $r = d = 0.3$ and $h_f = h_g = 1$.** The left panel shows the relationship-type classification for all 972 parameter combinations (concave: 65, convex: 338, linear: 59, and other: 510). The right panel shows the variable importance for classifying relationship types based on species interaction characteristics. Variable importance was quantified as the mean decrease in prediction accuracy after randomly permuting each predictor. The Y-axis represents the mean decrease in prediction accuracy. Error bars indicate 95% confidence intervals (CIs). The mean classification accuracy was 83.41 ± 0.09% (95% CIs).

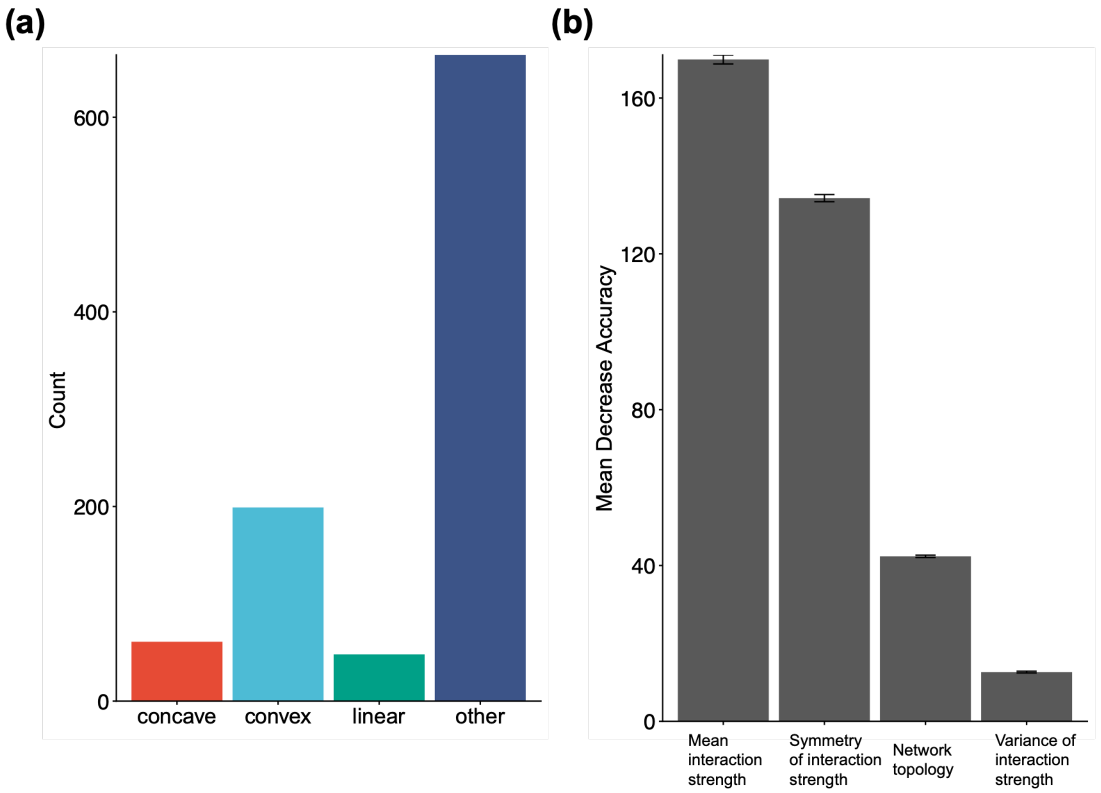


**Figure S14. Classification of increasing relationships between species pool size and the number of alternative stable states (ASSs) when $r = d = 0.3$ and $h_f = h_g = 10$.** The left panel shows the relationship-type classification for all 972 parameter combinations (concave: 61, convex: 199, linear: 48, and other: 664). The right panel shows the variable importance for classifying relationship types based on species interaction characteristics. Variable importance was quantified as the mean decrease in prediction accuracy after randomly permuting each predictor. The Y-axis represents the mean decrease in prediction accuracy. Error bars indicate 95% confidence intervals (CIs). The mean classification accuracy was 87.56 ± 0.09% (95% CIs).

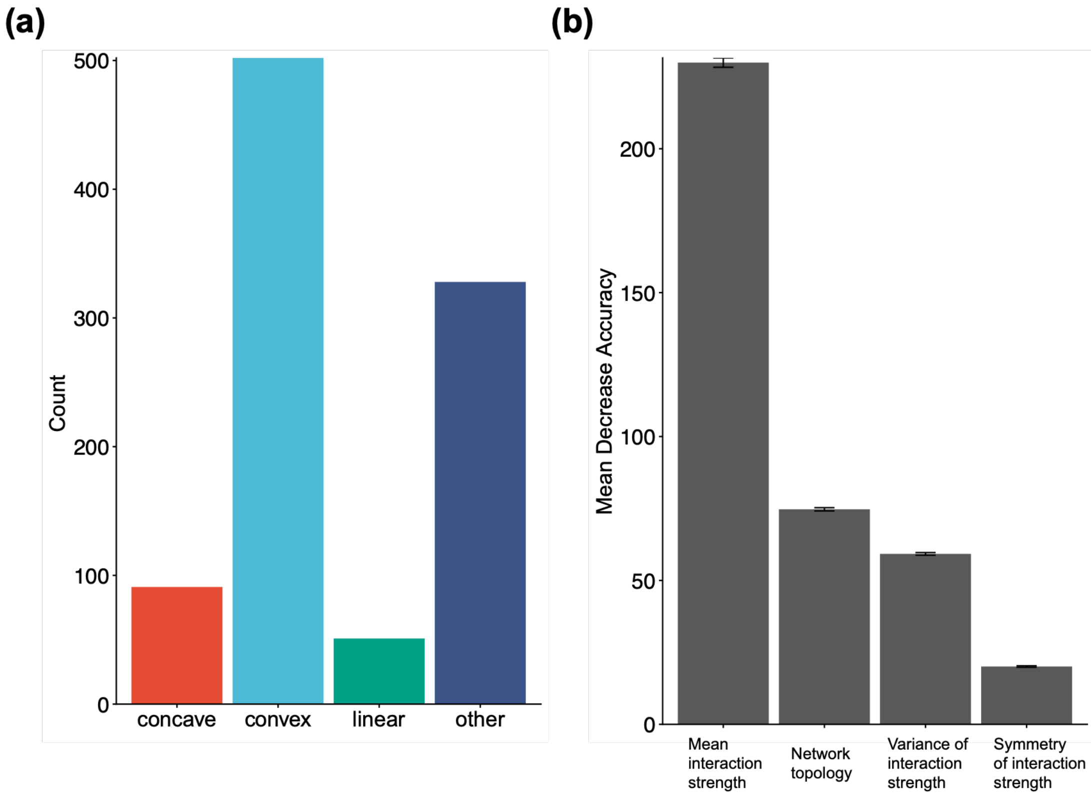


**Figure S15. Classification of increasing relationships between species pool size and the number of alternative stable states (ASSs) when $r = d = 0.6$ and $h_f = h_g = 0.1$.** The left panel shows the relationship-type classification for all 972 parameter combinations (concave: 91, convex: 502, linear: 51, and other: 328). The right panel shows the variable importance for classifying relationship types based on species interaction characteristics. Variable importance was quantified as the mean decrease in prediction accuracy after randomly permuting each predictor. The Y-axis represents the mean decrease in prediction accuracy. Error bars indicate 95% confidence intervals (CIs). The mean classification accuracy was 84.50 ± 0.08% (95% CIs).

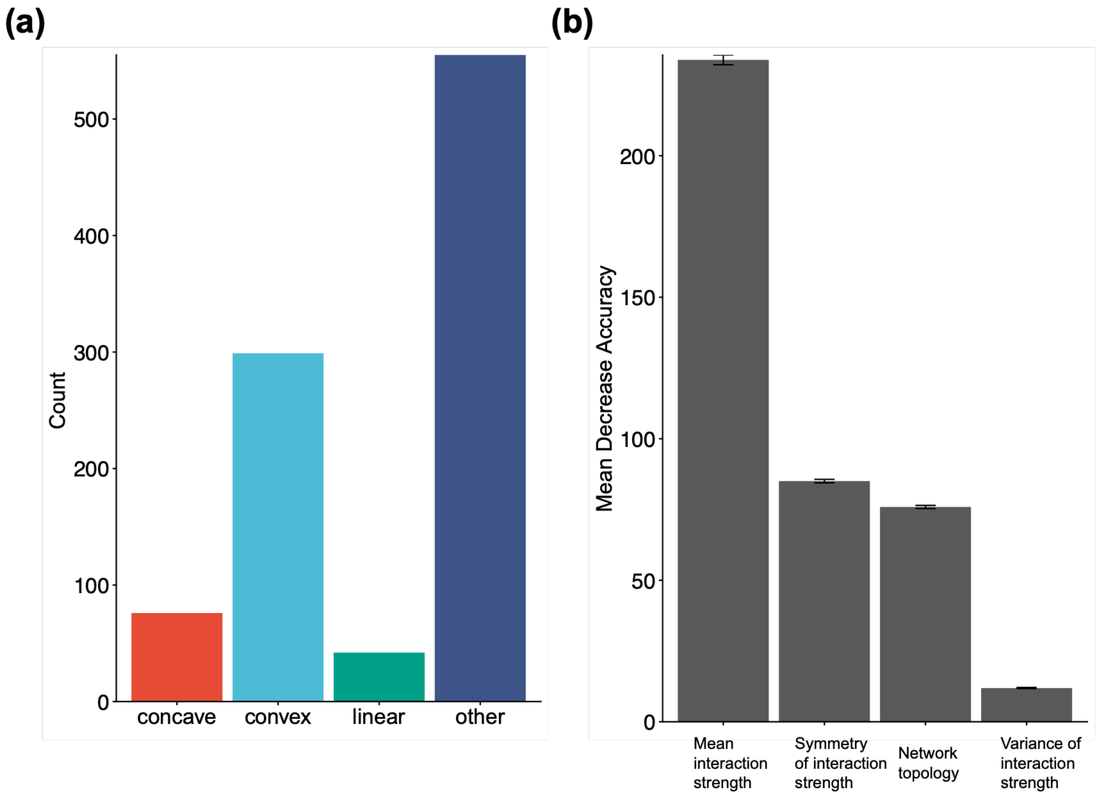


**Figure S16. Classification of increasing relationships between species pool size and the number of alternative stable states (ASSs) when $r = d = 0.6$ and $h_f = h_g = 1$.** The left panel shows the relationship-type classification for all 972 parameter combinations (concave: 76, convex: 299, linear: 42, and other: 555). The right panel shows the variable importance for classifying relationship types based on species interaction characteristics. Variable importance was quantified as the mean decrease in prediction accuracy after randomly permuting each predictor. The Y-axis represents the mean decrease in prediction accuracy. Error bars indicate 95% confidence intervals (CIs). The mean classification accuracy was 86.33 ± 0.08% (95% CIs).

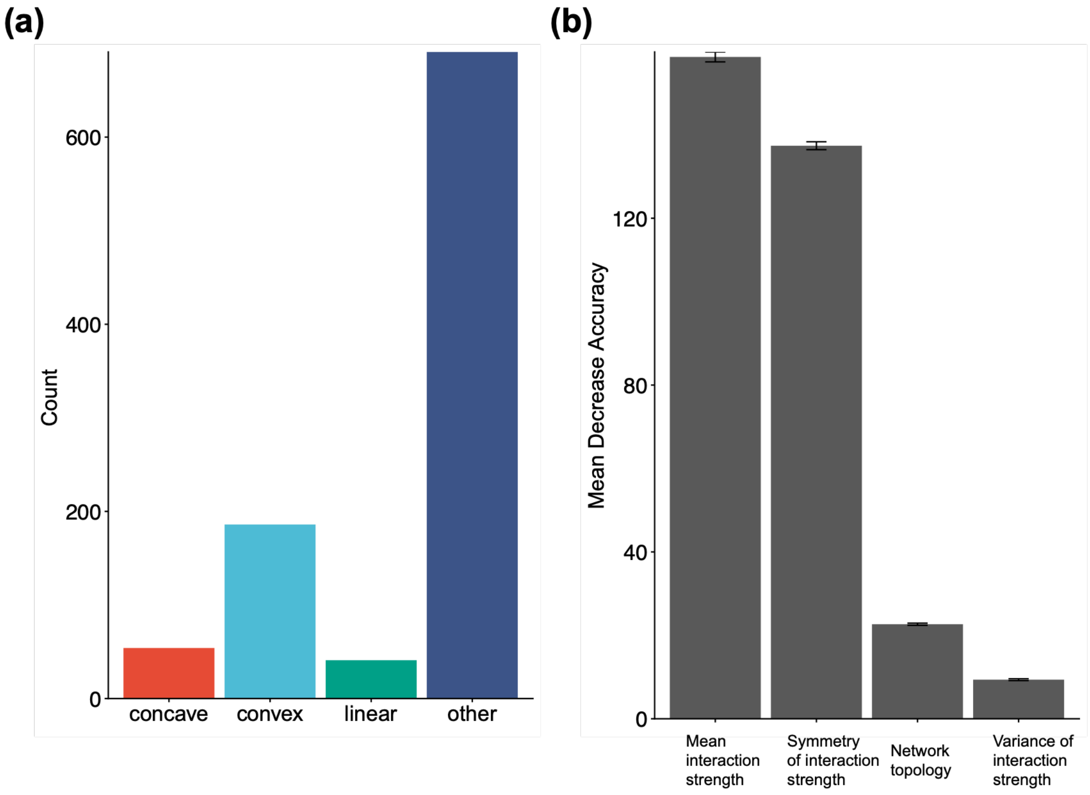


**Figure S17. Classification of increasing relationships between species pool size and the number of alternative stable states (ASSs) when $r = d = 0.6$ and $h_f = h_g = 10$.** The left panel shows the relationship-type classification for all 972 parameter combinations (concave: 54, convex: 186, linear: 41, and other: 691). The right panel shows the variable importance for classifying relationship types based on species interaction characteristics. Variable importance was quantified as the mean decrease in prediction accuracy after randomly permuting each predictor. The Y-axis represents the mean decrease in prediction accuracy. Error bars indicate 95% confidence intervals (CIs). The mean classification accuracy was 86.20 ± 0.08% (95% CIs).

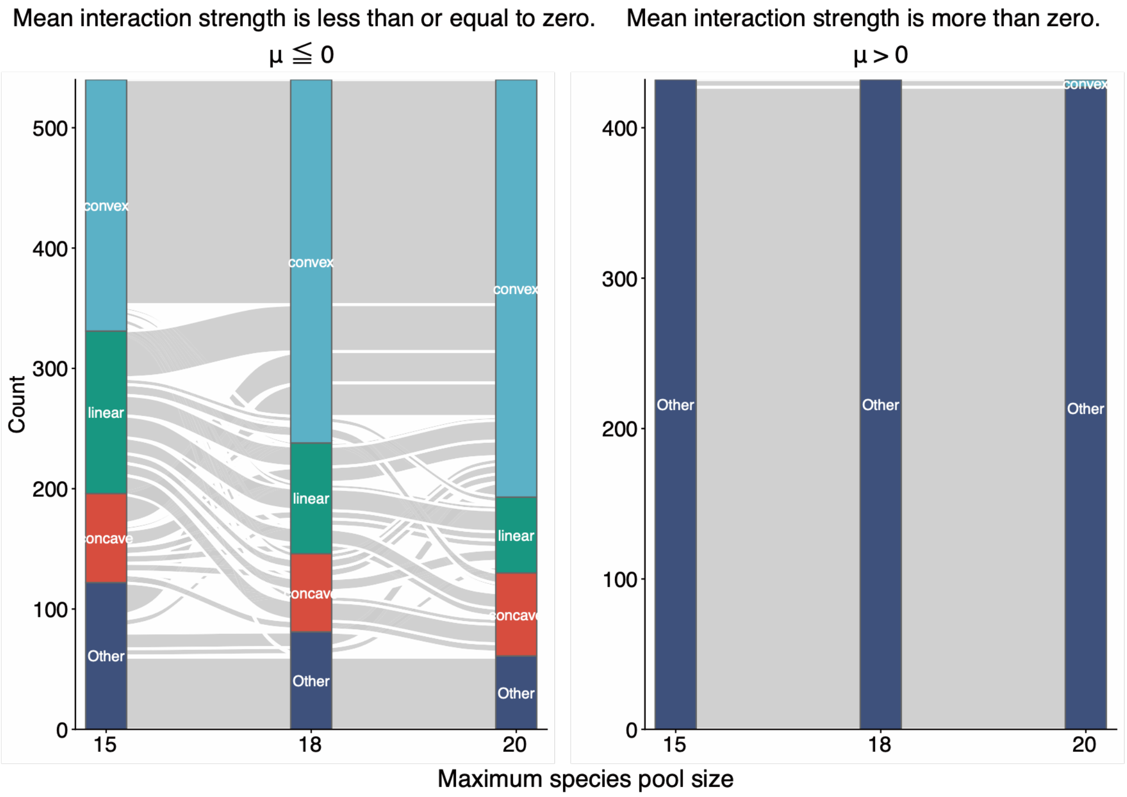


**Figure S18. Changes in relationship-type classifications with maximum species pool size.** The figure shows transitions in relationship-type classifications as the maximum species pool size increases from 15 to 20 species. Flows indicate transitions between relationship types. The left panel shows parameter combinations with mean interaction strength ($\mu$) less than or equal to 0, whereas the right panel shows those with mean interaction strength greater than 0. Increasing the maximum species pool size increased the proportion of relationships classified as "convex" when mean interaction strength was less than or equal to 0.

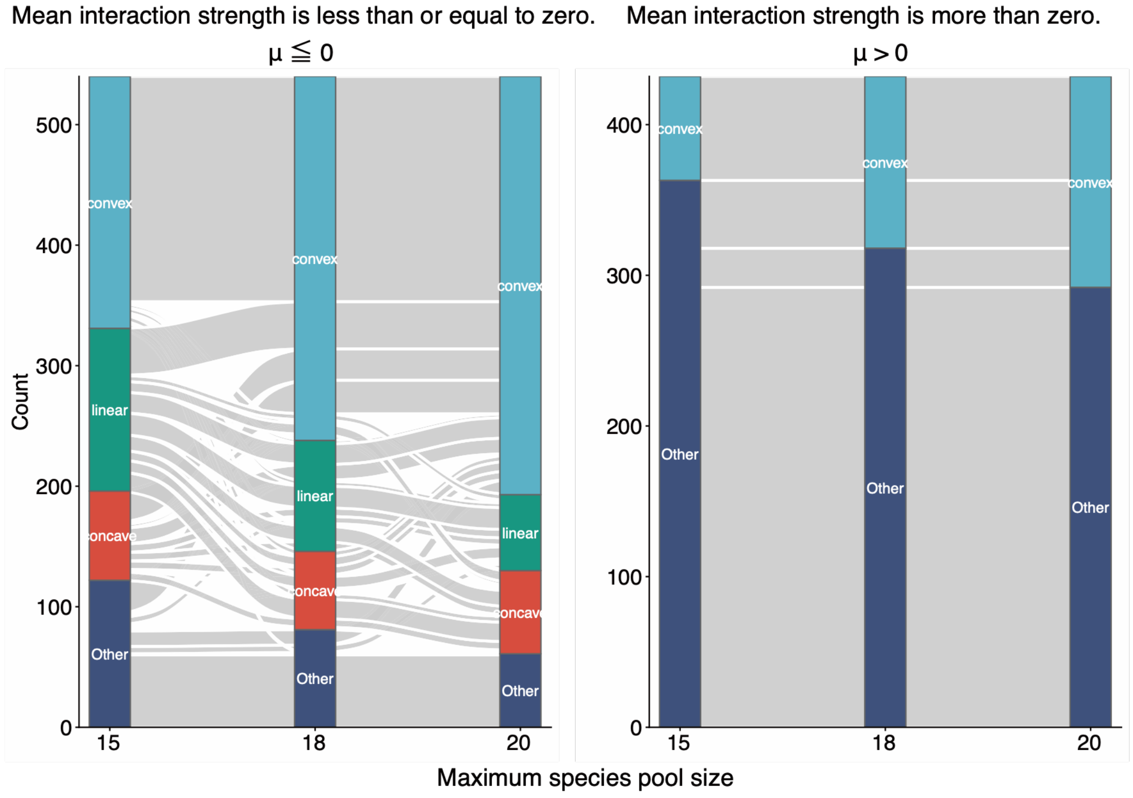


**Figure S19. Changes in relationship-type classifications with maximum species pool size when $r = d = 0.1$ and $h_f = h_g = 0.1$.** The figure shows transitions in relationship-type classifications as the maximum species pool size increases from 15 to 20 species. Flows indicate transitions between relationship types. The left panel shows parameter combinations with mean interaction strength ($\mu$) less than or equal to 0, whereas the right panel shows those with mean interaction strength greater than 0.

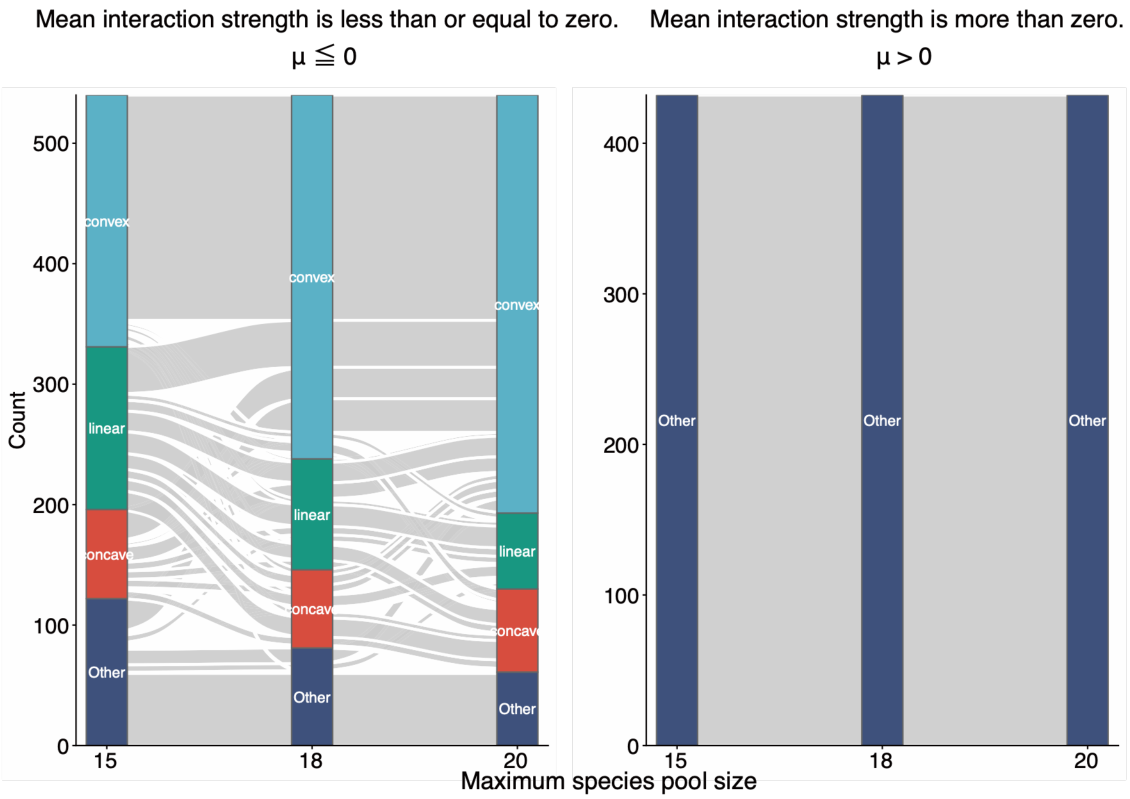


**Figure S20. Changes in relationship-type classifications with maximum species pool size when $r = d = 0.1$ and $h_f = h_g = 10$.** The figure shows transitions in relationship-type classifications as the maximum species pool size increases from 15 to 20 species. Flows indicate transitions between relationship types. The left panel shows parameter combinations with mean interaction strength ($\mu$) less than or equal to 0, whereas the right panel shows those with mean interaction strength greater than 0.

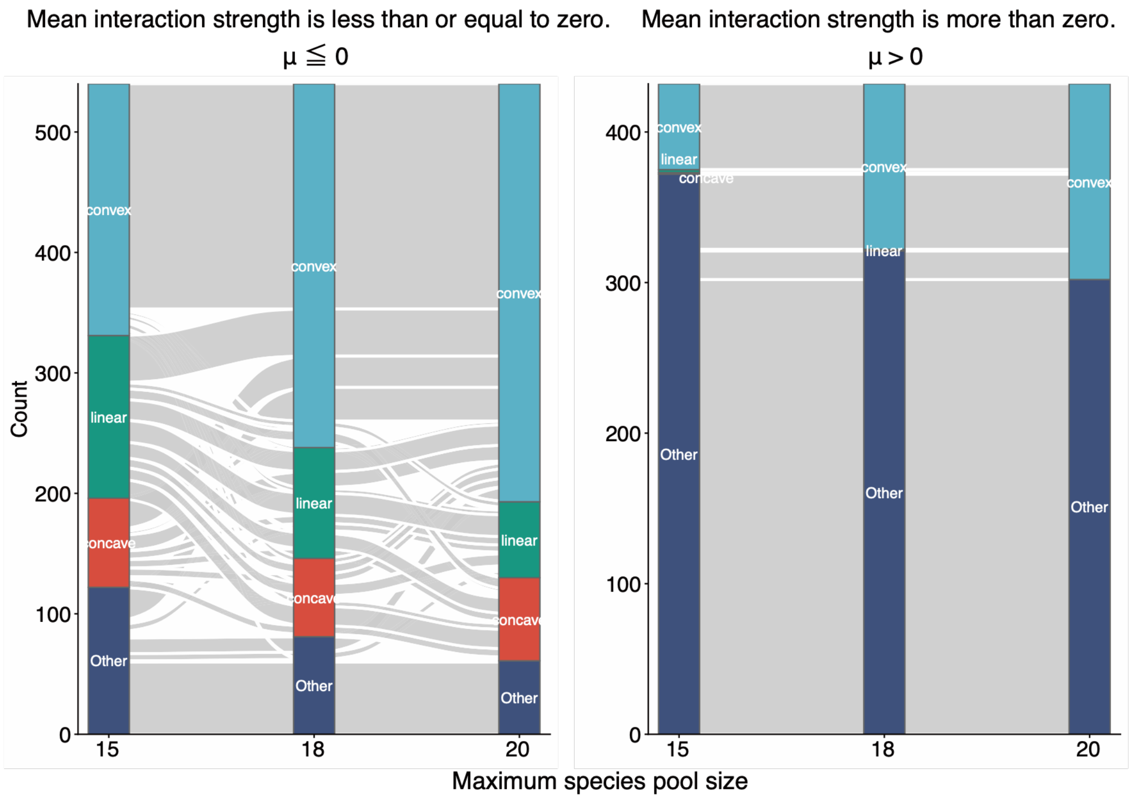


**Figure S21. Changes in relationship-type classifications with maximum species pool size when $r$ = $d$ = 0.3 and $h_f$ = $h_g$ = 0.1.** The figure shows transitions in relationship-type classifications as the maximum species pool size increases from 15 to 20 species. Flows indicate transitions between relationship types. The left panel shows parameter combinations with mean interaction strength ($\mu$) less than or equal to 0, whereas the right panel shows those with mean interaction strength greater than 0.

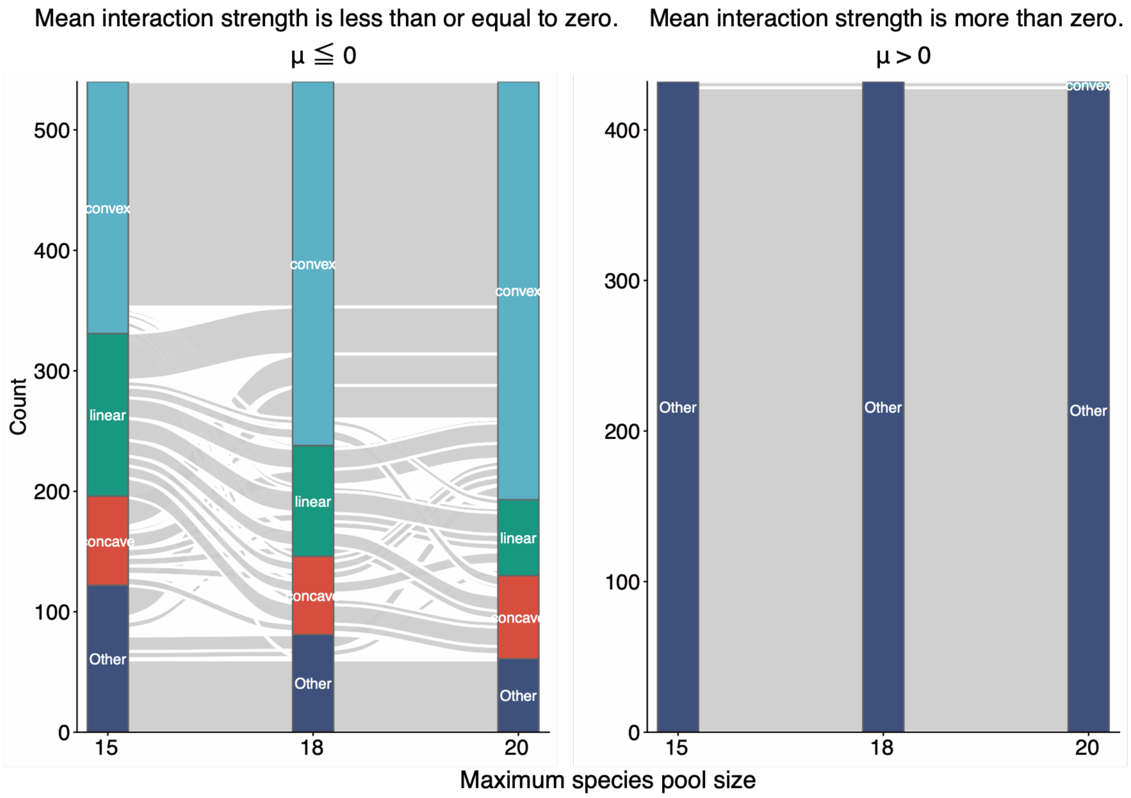


**Figure S22. Changes in relationship-type classifications with maximum species pool size when $r = d = 0.3$ and $h_f = h_g = 1$.** The figure shows transitions in relationship-type classifications as the maximum species pool size increases from 15 to 20 species. Flows indicate transitions between relationship types. The left panel shows parameter combinations with mean interaction strength ($\mu$) less than or equal to 0, whereas the right panel shows those with mean interaction strength greater than 0.

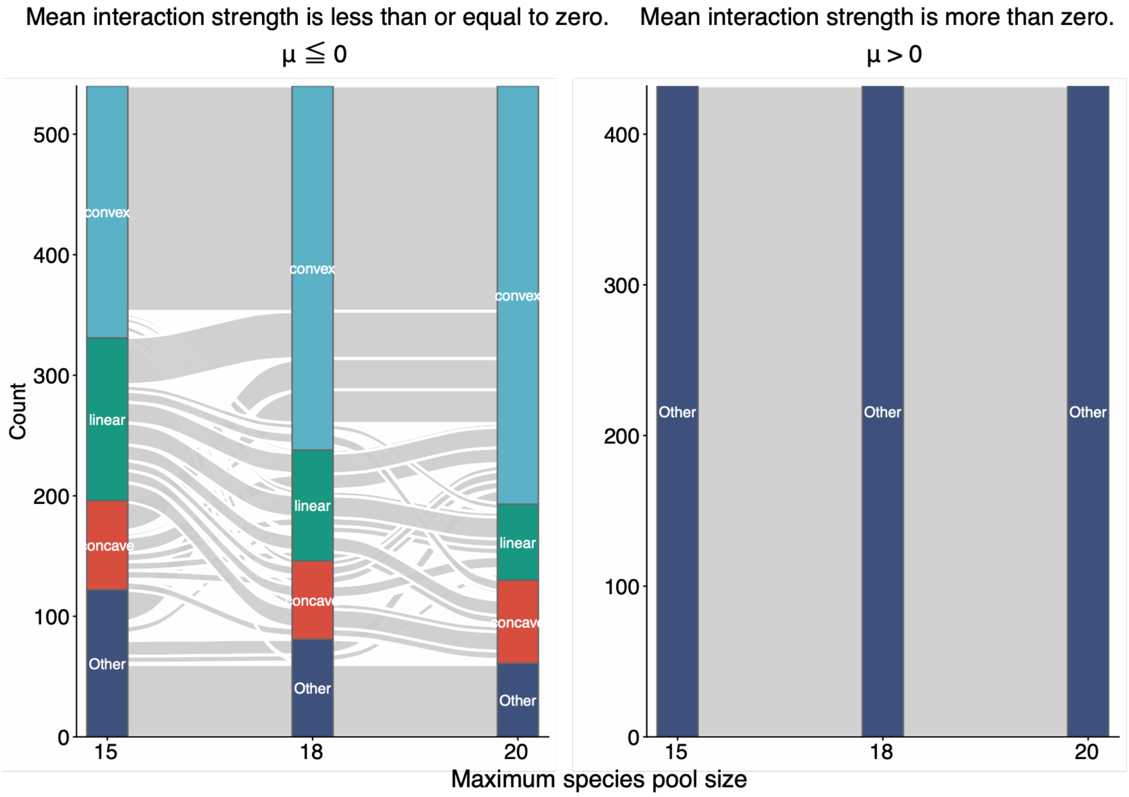


**Figure S23. Changes in relationship-type classifications with maximum species pool size when $r = d = 0.3$ and $h_f = h_g = 10$.** The figure shows transitions in relationship-type classifications as the maximum species pool size increases from 15 to 20 species. Flows indicate transitions between relationship types. The left panel shows parameter combinations with mean interaction strength ($\mu$) less than or equal to 0, whereas the right panel shows those with mean interaction strength greater than 0.

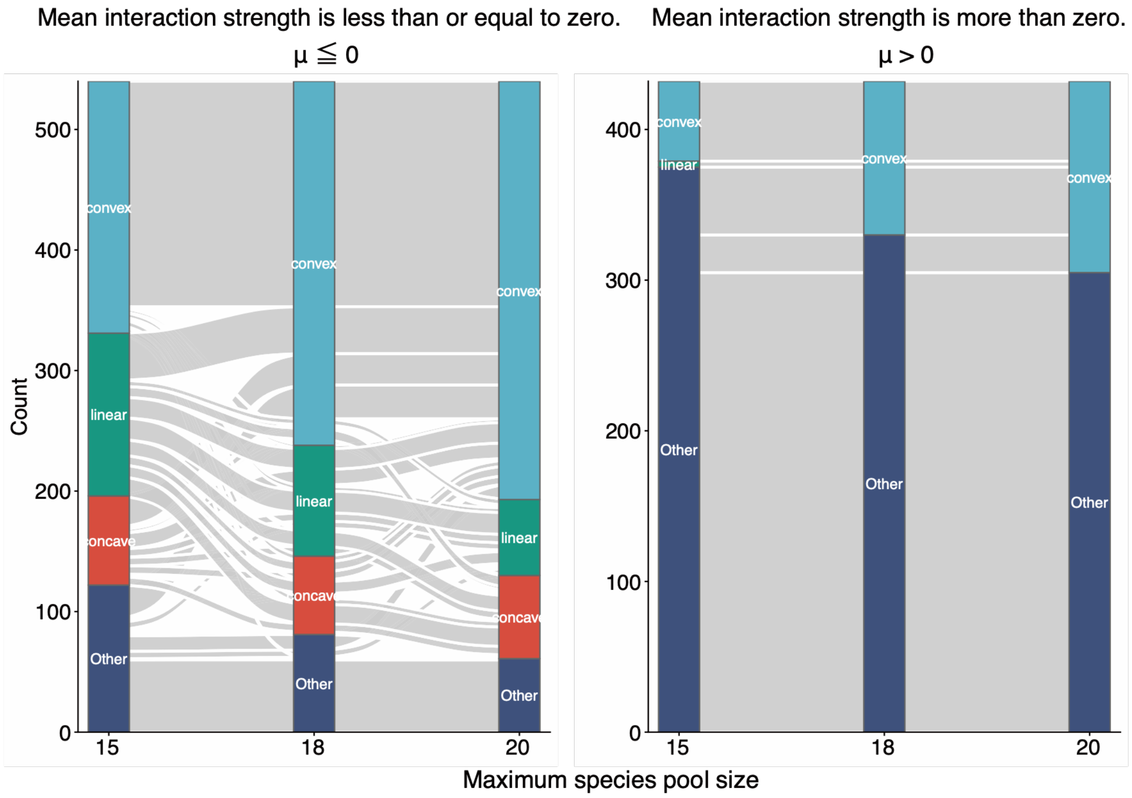


**Figure S24. Changes in relationship-type classifications with maximum species pool size when $r = d = 0.6$ and $h_f = h_g = 0.1$.** The figure shows transitions in relationship-type classifications as the maximum species pool size increases from 15 to 20 species. Flows indicate transitions between relationship types. The left panel shows parameter combinations with mean interaction strength ($\mu$) less than or equal to 0, whereas the right panel shows those with mean interaction strength greater than 0.

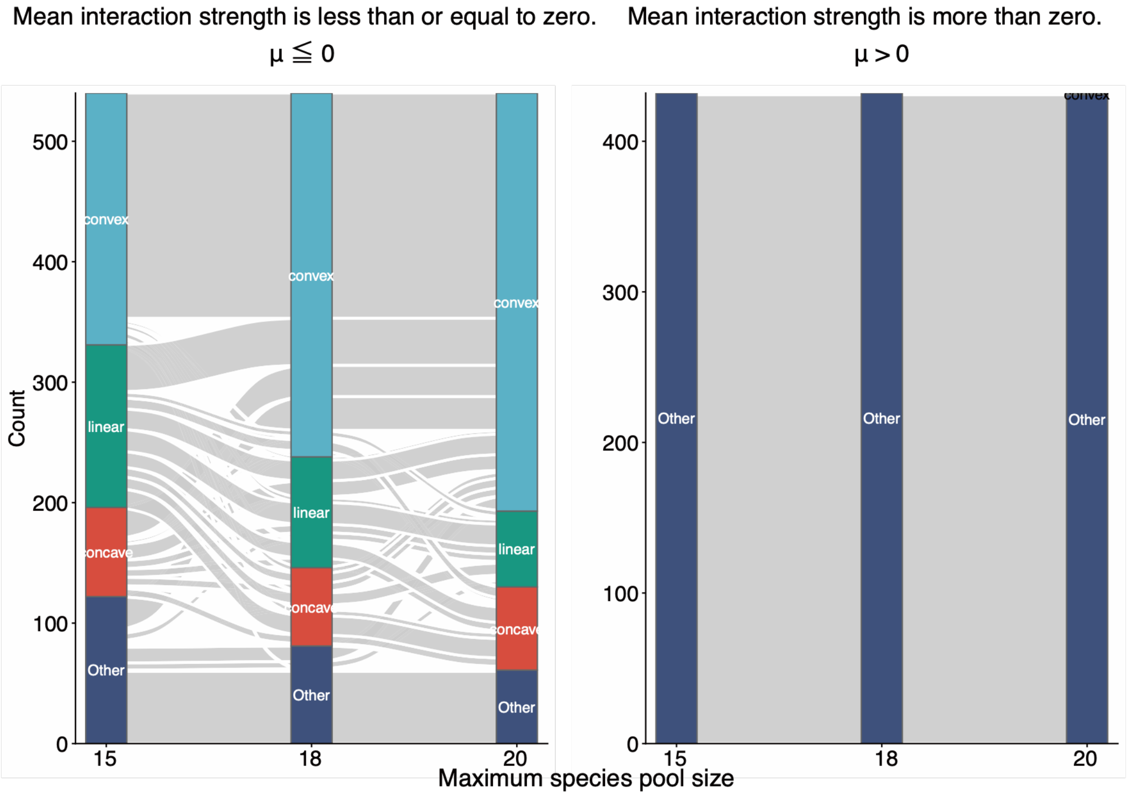


**Figure S25. Changes in relationship-type classifications with maximum species pool size when $r = d = 0.6$ and $h_f = h_g = 1$.** The figure shows transitions in relationship-type classifications as the maximum species pool size increases from 15 to 20 species. Flows indicate transitions between relationship types. The left panel shows parameter combinations with mean interaction strength ($\mu$) less than or equal to 0, whereas the right panel shows those with mean interaction strength greater than 0.

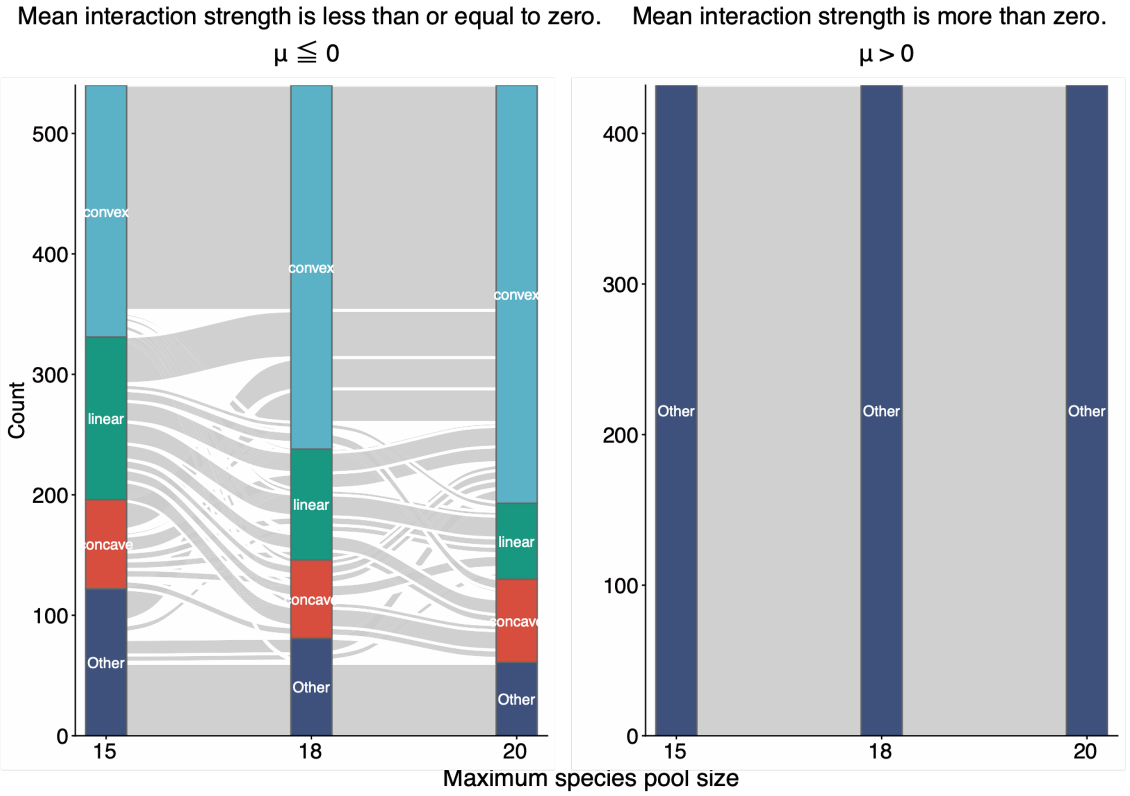


**Figure S26. Changes in relationship-type classifications with maximum species pool size when $r = d = 0.6$ and $h_f = h_g = 10$.** The figure shows transitions in relationship-type classifications as the maximum species pool size increases from 15 to 20 species. Flows indicate transitions between relationship types. The left panel shows parameter combinations with mean interaction strength ($\mu$) less than or equal to 0, whereas the right panel shows those with mean interaction strength greater than 0.

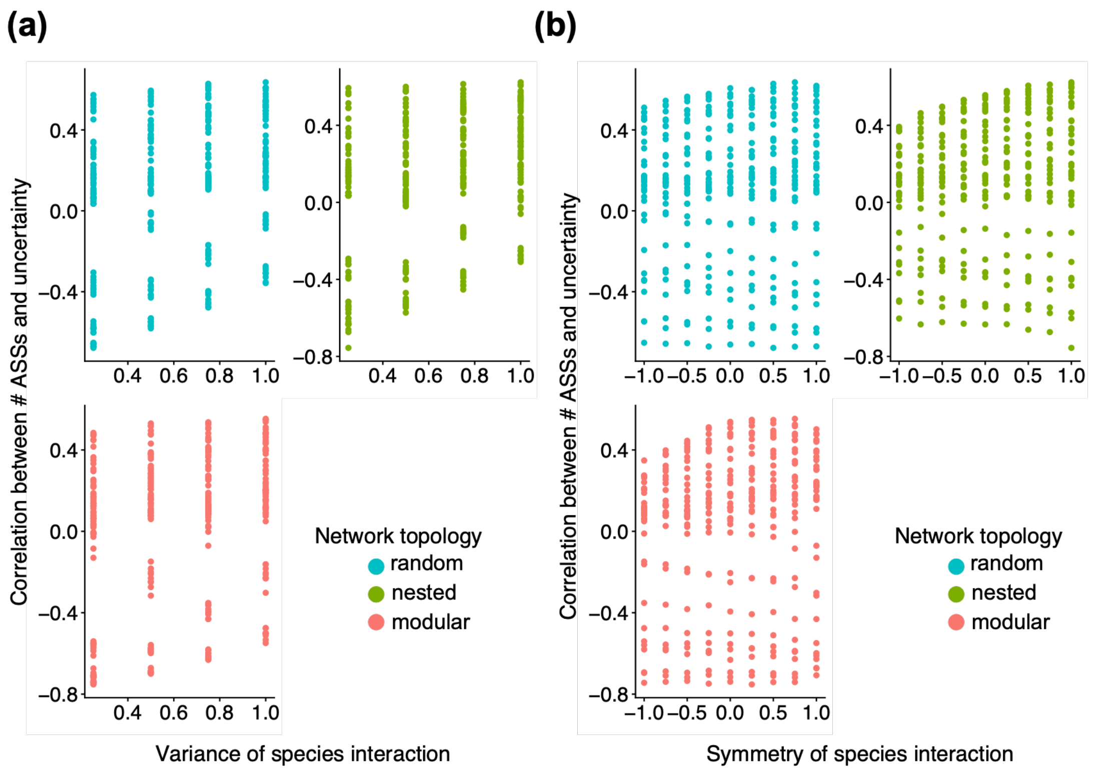


**Figure S27. Relationships between species interaction properties and Kendall's rank correlation coefficient ($\tau$) between the number of alternative stable states (ASSs) and community uncertainty.** (a) Relationship between the variance of interaction strengths ($\sigma^2$) and Kendall's $\tau$. No clear association was observed (Pearson's correlation = −0.29). (b) Relationship between interaction symmetry ($\rho$) and Kendall's $\tau$. No clear association was observed (Pearson's correlation = −0.06). Colors indicate network topology (blue: random, green: nested, and red: modular).

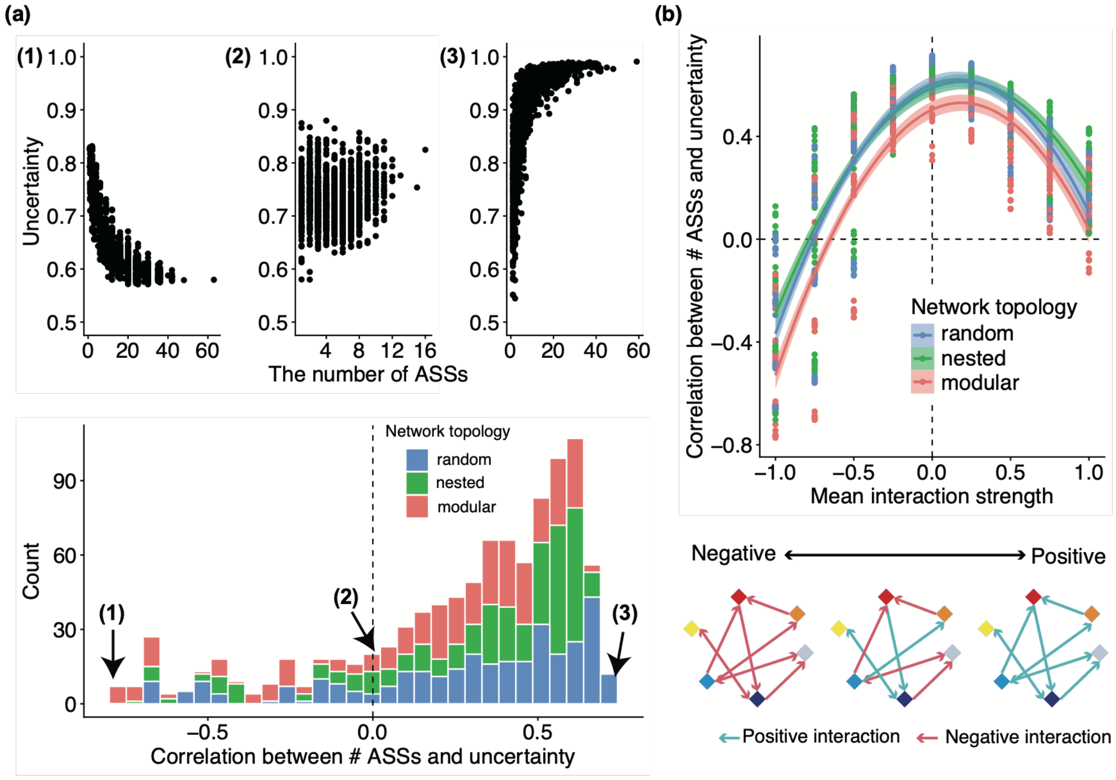


**Figure S28. Relationship between the number of alternative stable states (ASSs) and community uncertainty when $r = d = 0.1$ and $h_f = h_g = 0.1$.** (a) Kendall's rank correlation coefficient ($\tau$) ranged from −0.72 to 0.77. The upper panels show representative examples of the relationship between the number of ASSs and community uncertainty for the parameter combinations yielding the smallest (left; $\mu = -1,\ \sigma^2 = 0.25, \rho = -0.5$, modular network), near-zero (center; $\mu = -1,\ \sigma^2 = 1, \rho = 0.25$, random network), and largest (right; $\mu = 0$, $\sigma^2 = 0.75, \rho = 1$, random network) values of Kendall's $\tau$. The lower panel shows the distribution of Kendall's $\tau$ across all parameter combinations. Colors indicate network topology. The dashed vertical line indicates $\tau = 0$. (b) Relationship between mean interaction strength ($\mu$) and Kendall's $\tau$. Solid lines show quadratic regressions and shaded areas indicate 95% confidence intervals (CIs). Colors denote network topology. The dashed horizontal line indicates $\tau = 0$, and the dashed vertical line indicates $\mu = 0$. Pearson's correlation coefficients between Kendall's $\tau$ and interaction variance and interaction symmetry were −0.37 and 0.07, respectively.

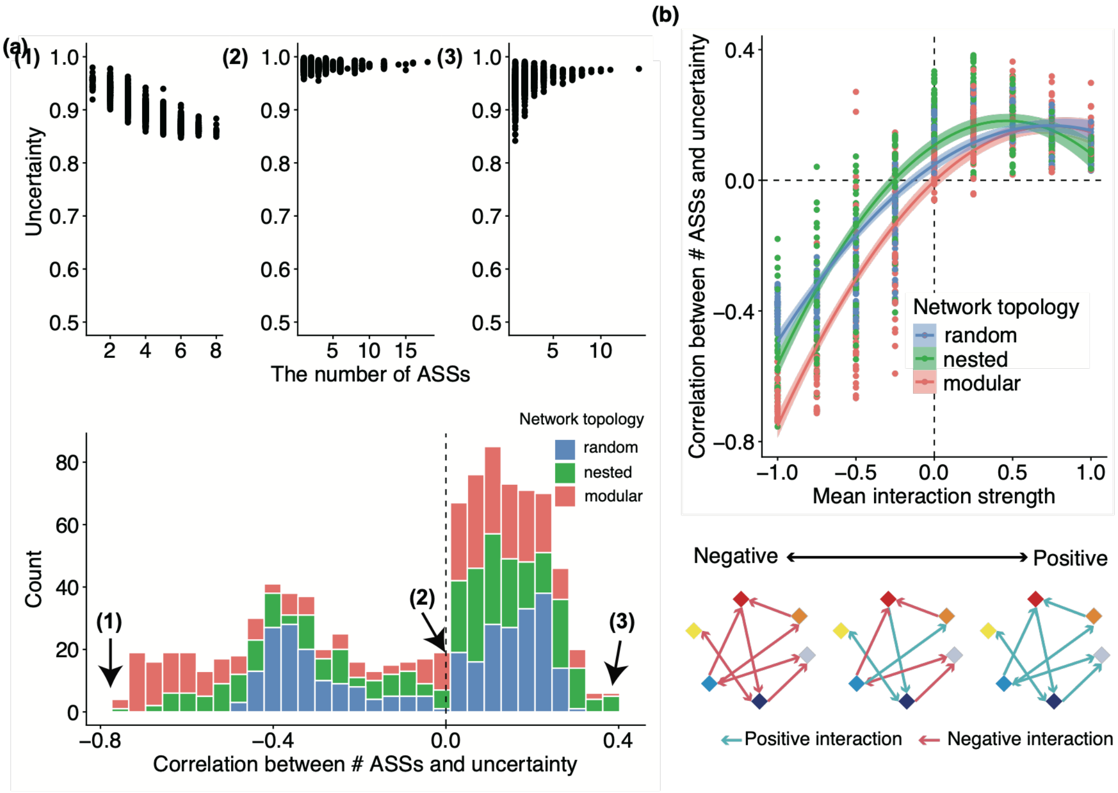


**Figure S29. Relationship between the number of alternative stable states (ASSs) and community uncertainty when $r = d = 0.1$ and $h_f = h_g = 10$.** (a) Kendall's rank correlation coefficient ($\tau$) ranged from −0.38 to 0.75. The upper panels show representative examples of the relationship between the number of ASSs and community uncertainty for the parameter combinations yielding the smallest (left; $\mu = -1,\ \sigma^2 = 0.25, \rho = 1$, nested network), near-zero (center; $\mu = 0,\ \sigma^2 = 0.25, \rho = -0.25$, modular network), and largest (right; $\mu = 0.25$, $\sigma^2 = 0.75, \rho = 1$, nested network) values of Kendall's $\tau$. The lower panel shows the distribution of Kendall's $\tau$ across all parameter combinations. Colors indicate network topology. The dashed vertical line indicates $\tau = 0$. (b) Relationship between mean interaction strength ($\mu$) and Kendall's $\tau$. Solid lines show quadratic regressions and shaded areas indicate 95% confidence intervals (CIs). Colors denote network topology. The dashed horizontal line indicates $\tau = 0$, and the dashed vertical line indicates $\mu = 0$. Pearson's correlation coefficients between Kendall's $\tau$ and interaction variance and interaction symmetry were −0.21 and 0.07, respectively.

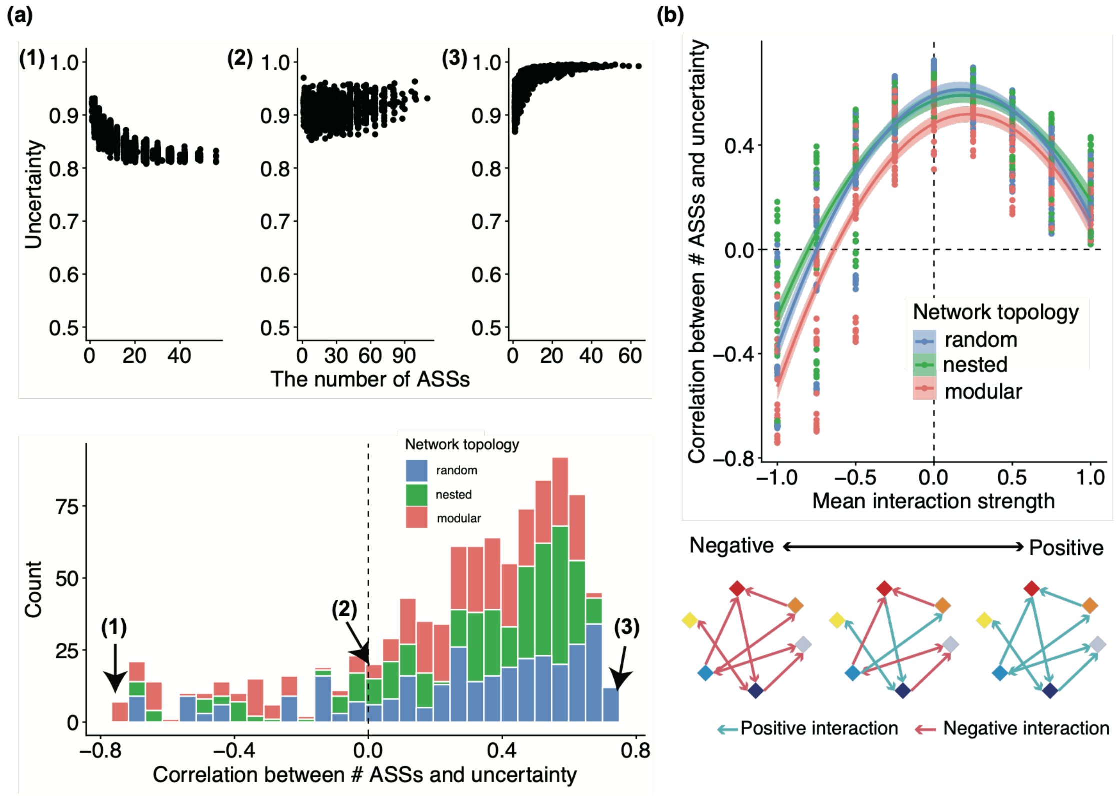


**Figure S30. Relationship between the number of alternative stable states (ASSs) and community uncertainty when $r = d = 0.3$ and $h_f = h_g = 0.1$.** (a) Kendall's rank correlation coefficient ($\tau$) ranged from −0.72 to 0.74. The upper panels show representative examples of the relationship between the number of ASSs and community uncertainty for the parameter combinations yielding the smallest (left; $\mu = -1$, $\sigma^2 = 0.25, \rho = -0.25$, modular network), near-zero (center; $\mu = -0.75$, $\sigma^2 = 0.75, \rho = 1$, modular network), and largest (right; $\mu = 0$, $\sigma^2 = 1, \rho = 1$, random network) values of Kendall's $\tau$. The lower panel shows the distribution of Kendall's $\tau$ across all parameter combinations. Colors indicate network topology. The dashed vertical line indicates $\tau = 0$. (b) Relationship between mean interaction strength ($\mu$) and Kendall's $\tau$. Solid lines show quadratic regressions and shaded areas indicate 95% confidence intervals (CIs). Colors denote network topology. The dashed horizontal line indicates $\tau = 0$, and the dashed vertical line indicates $\mu = 0$. Pearson's correlation coefficients between Kendall's $\tau$ and interaction variance and interaction symmetry were −0.38 and 0.08, respectively.

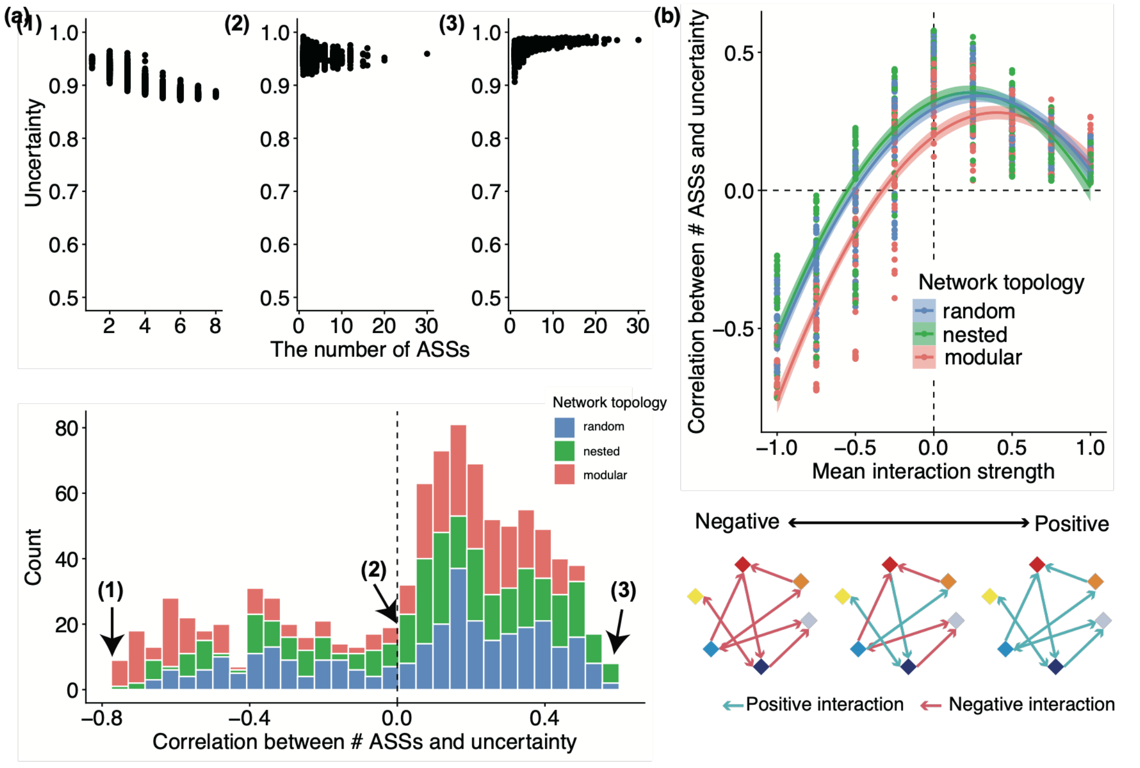


**Figure S31. Relationship between the number of alternative stable states (ASSs) and community uncertainty when $r = d = 0.3$ and $h_f = h_g = 1$.** (a) Kendall's rank correlation coefficient ($\tau$) ranged from −0.58 to 0.75. The upper panels show representative examples of the relationship between the number of ASSs and community uncertainty for the parameter combinations yielding the smallest (left; $\mu = -1,\ \sigma^2 = 0.25, \rho = 1$, nested network), near-zero (center; $\mu = -0.5,\ \sigma^2 = 1, \rho = -0.5$, modular network), and largest (right; $\mu = 0,\ \sigma^2 = 1, \rho = 0.75$, nested network) values of Kendall's $\tau$. The lower panel shows the distribution of Kendall's $\tau$ across all parameter combinations. Colors indicate network topology. The dashed vertical line indicates $\tau = 0$. (b) Relationship between mean interaction strength ($\mu$) and Kendall's $\tau$. Solid lines show quadratic regressions and shaded areas indicate 95% confidence intervals (CIs). Colors denote network topology. The dashed horizontal line indicates $\tau = 0$, and the dashed vertical line indicates $\mu = 0$. Pearson's correlation coefficients between Kendall's $\tau$ and interaction variance and interaction symmetry were −0.30 and −0.05, respectively.

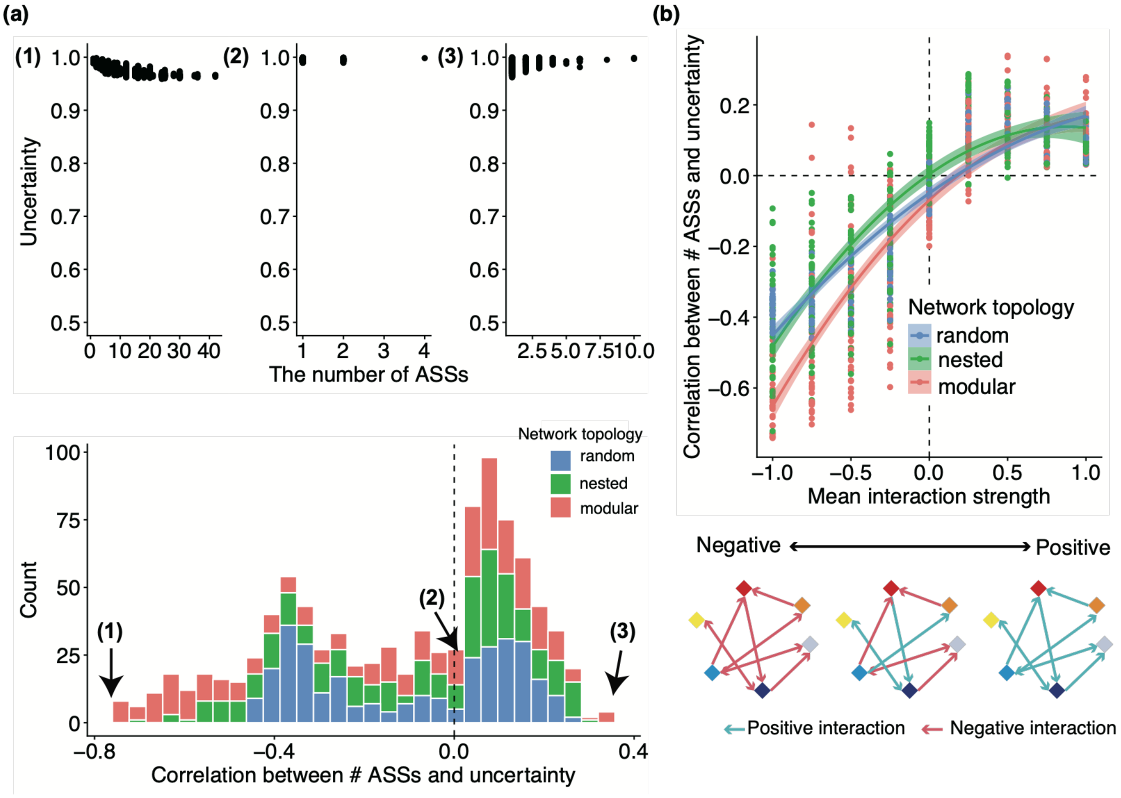


**Figure S32. Relationship between the number of alternative stable states (ASSs) and community uncertainty when $r = d = 0.3$ and $h_f = h_g = 10$.** (a) Kendall's rank correlation coefficient ($\tau$) ranged from −0.34 to 0.74. The upper panels show representative examples of the relationship between the number of ASSs and community uncertainty for the parameter combinations yielding the smallest (left; $\mu = -1$, $\sigma^2 = 0.25, \rho = 0.25$, modular network), near-zero (center; $\mu = 0.25$, $\sigma^2 = 0.5, \rho = -1$, modular network), and largest (right; $\mu = 0.5$, $\sigma^2 = 0.75, \rho = 1$, modular network) values of Kendall's $\tau$. The lower panel shows the distribution of Kendall's $\tau$ across all parameter combinations. Colors indicate network topology. The dashed vertical line indicates $\tau = 0$. (b) Relationship between mean interaction strength ($\mu$) and Kendall's $\tau$. Solid lines show quadratic regressions and shaded areas indicate 95% confidence intervals (CIs). Colors denote network topology. The dashed horizontal line indicates $\tau = 0$, and the dashed vertical line indicates $\mu = 0$. Pearson's correlation coefficients between Kendall's $\tau$ and interaction variance and interaction symmetry were −0.17 and 0.13, respectively.

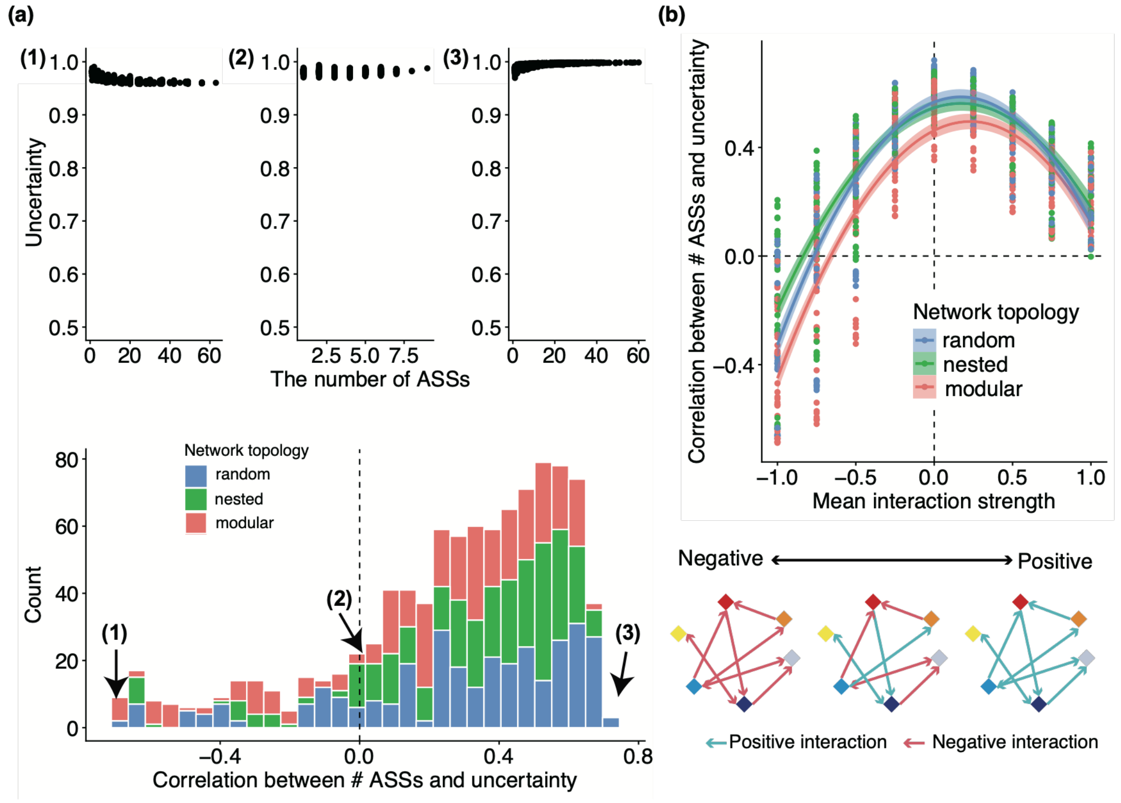


**Figure S33. Relationship between the number of alternative stable states (ASSs) and community uncertainty when $r = d = 0.6$ and $h_f = h_g = 0.1$.** (a) Kendall's rank correlation coefficient ($\tau$) ranged from −0.72 to 0.69. The upper panels show representative examples of the relationship between the number of ASSs and community uncertainty for the parameter combinations yielding the smallest (left; $\mu = -1$, $\sigma^2 = 0.25, \rho = 0$, modular network), near-zero (center; $\mu = -0.75$, $\sigma^2 = 0.5, \rho = -0.25$, nested network), and largest (right; $\mu = 0$, $\sigma^2 = 1, \rho = 1$, random network) values of Kendall's $\tau$. The lower panel shows the distribution of Kendall's $\tau$ across all parameter combinations. Colors indicate network topology. The dashed vertical line indicates $\tau = 0$. (b) Relationship between mean interaction strength ($\mu$) and Kendall's $\tau$. Solid lines show quadratic regressions and shaded areas indicate 95% confidence intervals (CIs). Colors denote network topology. The dashed horizontal line indicates $\tau = 0$, and the dashed vertical line indicates $\mu = 0$. Pearson's correlation coefficients between Kendall's $\tau$ and interaction variance and interaction symmetry were −0.39 and 0.10, respectively.

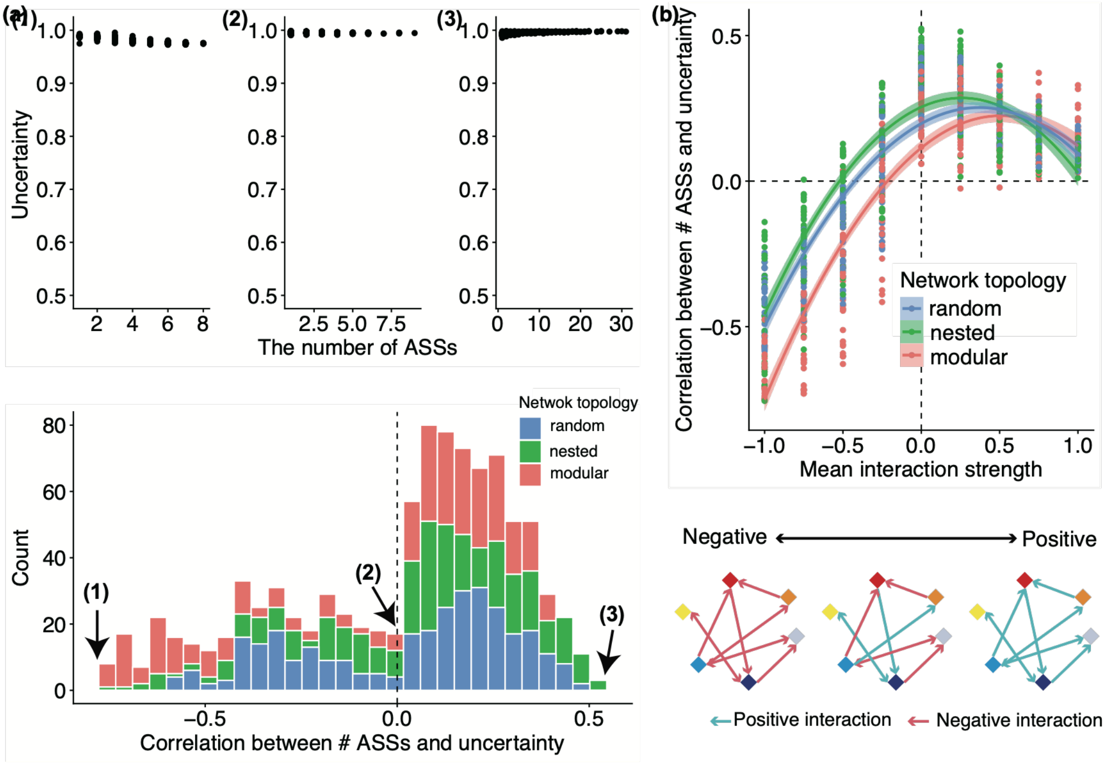


**Figure S34. Relationship between the number of alternative stable states (ASSs) and community uncertainty when $r = d = 0.6$ and $h_f = h_g = 1$.** (a) Kendall's rank correlation coefficient ($\tau$) ranged from −0.52 to 0.76. The upper panels show representative examples of the relationship between the number of ASSs and community uncertainty for the parameter combinations yielding the smallest (left; $\mu = -1$, $\sigma^2 = 0.25, \rho = 1$, nested network), near-zero (center; $\mu = -0.25$, $\sigma^2 = 0.25, \rho = -0.75$, nested network), and largest (right; $\mu = 0$, $\sigma^2 = 1, \rho = 0.75$, nested network) values of Kendall's $\tau$. The lower panel shows the distribution of Kendall's $\tau$ across all parameter combinations. Colors indicate network topology. The dashed vertical line indicates $\tau = 0$. (b) Relationship between mean interaction strength ($\mu$) and Kendall's $\tau$. Solid lines show quadratic regressions and shaded areas indicate 95% confidence intervals (CIs). Colors denote network topology. The dashed horizontal line indicates $\tau = 0$, and the dashed vertical line indicates $\mu = 0$. Pearson's correlation coefficients between Kendall's $\tau$ and interaction variance and interaction symmetry were −0.29 and −0.36, respectively.

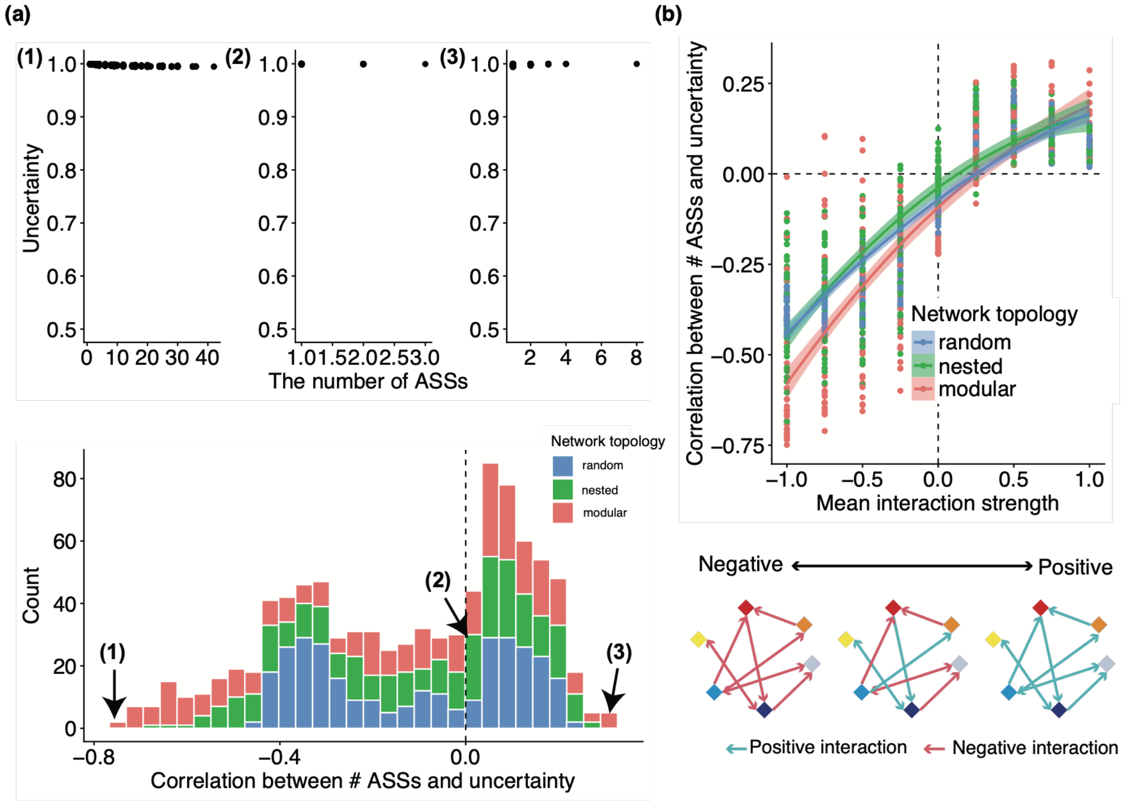


**Figure S35. Relationship between the number of alternative stable states (ASSs) and community uncertainty when $r = d = 0.6$ and $h_f = h_g = 10$.** (a) Kendall's rank correlation coefficient ($\tau$) ranged from −0.31 to 0.75. The upper panels show representative examples of the relationship between the number of ASSs and community uncertainty for the parameter combinations yielding the smallest (left; $\mu = -1,\ \sigma^2 = 0.25, \rho = 0.75$, modular network), near-zero (center; $\mu = 0, \sigma^2 = 1, \rho = -1$, nested network), and largest (right; $\mu = 0.75,\ \sigma^2 = 0.75, \rho = 1$, modular network) values of Kendall's $\tau$. The lower panel shows the distribution of Kendall's $\tau$ across all parameter combinations. Colors indicate network topology. The dashed vertical line indicates $\tau = 0$. (b) Relationship between mean interaction strength ($\mu$) and Kendall's $\tau$. Solid lines show quadratic regressions and shaded areas indicate 95% confidence intervals (CIs). Colors denote network topology. The dashed horizontal line indicates $\tau = 0$, and the dashed vertical line indicates $\mu = 0$. Pearson's correlation coefficients between Kendall's $\tau$ and interaction variance and interaction symmetry were −0.14 and 0.15, respectively.
